# The innate immunity protein C1QBP functions as a negative regulator of circulative transmission of *Potato leafroll virus* by aphids

**DOI:** 10.1101/2020.12.04.412668

**Authors:** Stacy L. DeBlasio, Jennifer Wilson, Cecilia Tamborindeguy, Richard S. Johnson, Patricia V. Pinheiro, Michael J. MacCoss, Stewart M. Gray, Michelle Heck

## Abstract

The vast majority of plant viruses are transmitted by insect vectors with many crucial aspects of the transmission process being mediated by key protein-protein interactions. Yet, very few vector proteins interacting with virus have been identified and functionally characterized. *Potato leafroll virus* (PLRV) is transmitted most effectively by *Myzus persicae*, the green peach aphid, in a circulative, non-propagative manner. Using an affinity purification strategy coupled to high-resolution mass spectrometry (AP-MS), we identified 11 proteins from *M. persicae* displaying high probability of interaction with PLRV and an additional 23 vector proteins with medium confidence interaction scores. Two of these proteins were confirmed to directly interact with the structural proteins of PLRV and other luteovirid species via yeast two-hybrid with an additional vector protein displaying binding specificity. Immunolocalization of one of these direct PLRV-interacting proteins, an orthologue of the human innate immunity protein complement component 1 Q subcomponent-binding protein (C1QBP), shows that MpC1QBP partially co-localizes with PLRV within cytoplasmic puncta and along the periphery of aphid gut epithelial cells. Chemical inhibition of C1QBP in the aphid leads to increased PLRV acquisition and subsequently increased titer in inoculated plants, supporting the role of C1QBP as a negative regulator of PLRV accumulation in *M. persicae*. We hypothesize that the innate immune function of C1QBP is conserved in aphids and represents the first instance of aphids mounting an immune response to a non-propagative plant virus. This study presents the first use of AP-MS for the *in vivo* isolation of functionally relevant insect vector-virus protein complexes.

## IMPORTANCE

The control of vector-borne disease is recognized as one of the major agricultural and human health challenges of today. Despite the importance of insect vectors, very little is known about the vector proteins that regulate transmission of viruses, especially viruses infecting plants. In this research, we adapt an emerging technique used to isolate host-pathogen interactions to identify insect vector-virus protein complexes. Using the aphid-borne, plant-infecting virus *Potato leafroll viru*s, we identified several vector proteins interacting with this virus and go on to show that one of these proteins may be a part of the aphid immune system that limits the amount of virus in the insect. Identifying and understanding the role of this and other insect proteins may be the first step to developing new strategies to control these and other insect-borne viruses.

## INTRODUCTION

Transmitting over 100 different plant virus species, aphids are the most prolific insect vectors in agroecosystems (1). Transmission of viruses by insect vectors, such as aphids, can be grouped into two general modes: circulative and non-circulative transmission, depending on the length and nature of the association of the virus with insect tissues. Circulative viruses traffic through insect vector tissues through a poorly described process of transcytosis, whereas non-circulative viruses largely adhere to insect mouthparts and some to the foregut (2). Moreover, plant viruses may or may not be propagative, or replicate, in their insect vector tissues. One important group of viruses vectored by aphids are the luteovirids (family *Luteoviridae*), which are transmitted in a circulative, non-propagative mode. The luteovirid *Potato leafroll virus* (PLRV, genus *Polerovirus*) and its primary vector, the green peach aphid, *Myzus persicae* (Hemiptera: Aphididae) are problematic in potato growing regions of the world, particularly where insecticides are not used to control aphid populations. PLRV is one of many viruses, which lead to degeneracy of potato and economically significant crop loss. Aphid populations rapidly become resistant to insecticides and thus, novel approaches to manage aphid-borne viral diseases are critically needed.

Circulative transmission of PLRV requires the successive passage of the virus through several membrane barriers of the aphid, most notably the gut and accessory salivary glands. Electron micrographs have detailed the transmission of luteovirids on the cellular level (3, 4). In the aphid gut, association of virus particles with receptors on the apical plasma membrane of epithelial cells initiates clathrin-mediated endocytosis (5). The virions traffic through the endomembrane system of gut epithelial cells in tubular vesicles that eventually fuse with the basal cell membrane, a process known as transcytosis. Virions are then released into the hemocoel (6). This aspect of the transmission process is referred to as acquisition. Luteovirid species show different affinity for various regions of the gut; PLRV and other viruses in the genus *Polerovirus* are acquired through the posterior midgut (7) whereas those in the genus *Luteovirus* are preferentially acquired through for the hindgut. Once in the hemocoel, virions are hypothesized to diffuse until encountering the accessory salivary glands, where they adhere to the basal lamina and the process of transcytosis occurs again allowing virions to be released into the salivary duct and spit into the host plant along with salivary secretions during feeding (8). Each luteovirid species is only transmitted by one or a few aphid vector species (9).

However, across all plant virus-vector systems, fewer than a dozen vector proteins involved in transmission have been identified and functionally validated (10–16) (reviewed in (17)). It is not understood to what extent these protein interactions are conserved across different virus-vector systems. Virus receptor proteins expressed in the vector are an excellent example. The ephrin receptor protein was recently identified as a putative receptor for the luteovirids *Turnip yellows virus, Beet mild yellowing virus,* and *Cucurbit aphid borne yellows virus*, all in the genus *Polerovirus* and transmitted by the aphid *M. persicae* (18). However, a different protein, aminopeptidase-N, was shown to be a putative receptor of *Pea enation mosaic virus* in its primary vector, *Acrythosiphon pisum* (19). While there appears to be specificity in the cell-surface receptors for the luteovirids, it is possible that once virions enter cells the interactions may be more conserved. For instance, the protein cyclophilin B has been implicated in the transmission of both the luteovirid *Cereal yellow dwarf virus-RPV* (CYDV-RPV, genus *Polerovirus*) by the aphid *Schizaphis graminum* (20) and the begomovirus *Tomato yellow leaf curl virus* (TYLCV, family *Geminiviridae*) transmitted by the whitefly *Bemisia tabaci* (11).

This is in contrast to the noncirculative viruses, where the same cuticle protein, *Myzus persicae* cuticle protein 4 (also known as stylin-01), is involved in adhesion of both *Cucumber mosaic virus* (CMV, *Bromoviridae: Cucumovirus*) (13) and *Cauliflower mosaic virus* (CaMV, *Caulimoviridae: Caulimovirus*) (15) to aphid stylets, even though CMV binds the stylet directly and CaMV uses the virus helper proteins P2 and P3 (21). However, noncirculative viruses show less vector specificity than luteovirids and other circulative viruses, as dozens of aphid vector species can transmit the same noncirculative virus (22).

Identifying vector proteins involved in the transmission process remains technically challenging due in part to the small size of insect vectors, the lack of genomic resources for these vectors, the difficulty of extracting vector proteins (especially cuticle proteins important for non-circulative viruses) and the relatively low amount of virus present compared to relative levels in plants (especially for non-propagative viruses). Nevertheless, many *in vitro* approaches have been used to address this challenge including far-western virus overlays (12, 23–27) and yeast two-hybrid (13, 14, 28, 29). Others have looked for differentially expressed genes and proteins (30–34) or proteomic phenotyping of vector and non-vector insects within the same species (35–37). Once identified, functional analysis of these proteins is another hurdle as genome editing is not routine or trivial for aphids and other non-model organism insect vector species. The possible roles of these candidate proteins are myriad: some may be defense response proteins from the vector, while other proteins crucial for the virus to move through the vector might be involved in other physiological functions. Moreover, plant viruses have been shown to manipulate their vectors on many levels, from counter-defense on the molecular level (16) to manipulating vector behavior and physiology (reviewed in (38)). Identifying these vector proteins and the role they play in transmission could be the key to developing strategies to block transmission. The activity of defense proteins can be enhanced, proteins the virus needs to complete its passage through the insect can be downregulated or edited to no longer bind to virus, and interfering with the virus’ ability to manipulate vector behavior can slow or prevent vectors from finding infected plant hosts (discussed in (39)). Therefore, characterizing the role of vector proteins in the transmission process is of great importance to implement strategies to protect crops.

In this work, we used affinity purification coupled to high-resolution mass spectrometry (AP-MS) to identify PLRV-*M. persicae* protein complexes formed within aphid tissues. Previously, we applied this technique to capture and identify PLRV-plant protein complexes directly from virus-infected mesophyll and phloem tissue (40–42). We adapted this workflow to rapidly isolate PLRV-aphid protein complexes from viruliferous aphids. Using a variety of cellular and molecular approaches, we probed the functional role of complement component 1 Q subcomponent-binding protein (C1QBP), the most enriched virus-interacting aphid protein identified by AP-MS, in the transmission of PLRV by its insect vector. Collectively, our data provide evidence for the conserved role of this protein in animal immunity.

## RESULTS

### Extraction of polerovirus-vector complexes requires different lysis buffer conditions from that used for plants

To determine an optimal buffer to use for capturing PLRV-associated protein complexes from aphids, we used a far-western approach to compare four routinely used extraction buffer compositions (40, 43) for their ability to extract PLRV-interacting aphid (Fig.1A, Table 1.). A denaturing buffer containing 2.5% SDS was used as a positive control as the strong detergent would extract the vast majority of proteins from the aphid. Whole, non-viruliferous, *M. persicae* were cryogenically lysed in a Mixer Mill using an increased number of grinding cycles than that used for plant tissue (40). The resulting powder was split equally and solubilized in the same volume of the different extraction buffers. Visual inspection of centrifugation-cleared homogenates showed differences in pigmentation, suggesting that, to some degree, each buffer resulted in the extraction of a different profile of molecules from the same pool of cryogenically lysed, tissue (Fig. 1A, top panel). The CHAPS-based buffer, optimal for extracting membrane bound proteins (43), resulted in a bluish-green hued homogenate, while extraction with the HEPES or TBT-based buffers supplemented with the non-ionic detergents Triton-X 100 and Tween-20, respectively, resulted in a yellowish-green tinted homogenate. Extraction with the Tris-based buffer (Fig. 1A, top panel, TRIS) was a combination of bluish-green and the reddish pigmentation observed when the same pool of tissue was extracted with SDS.

**FIG 1.**
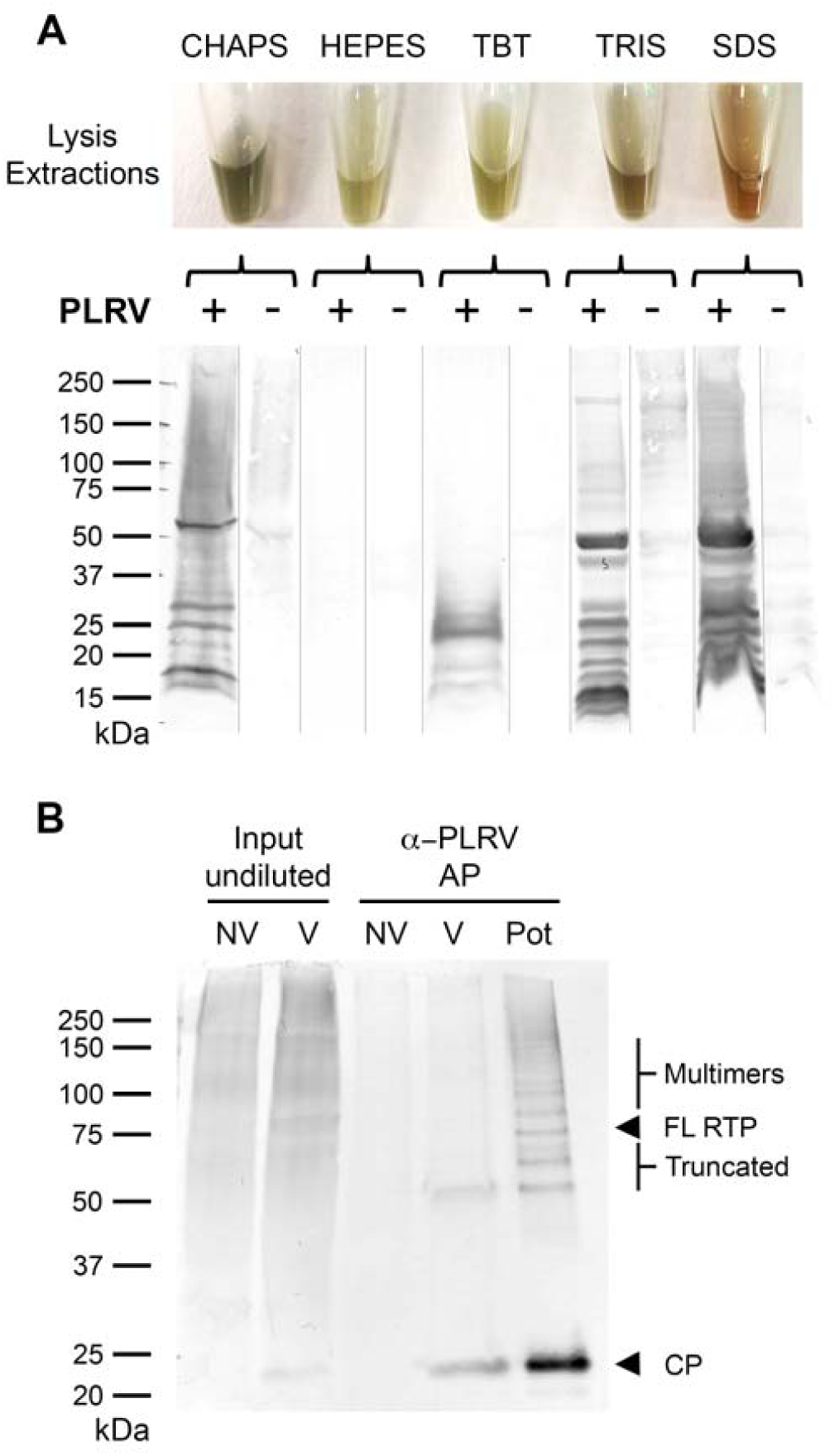
Comparison and validation of extraction conditions for isolating PLRV-vector protein complexes. (A) Four different affinity purification buffer compositions (CHAPS, HEPES, TBT and TRIS) were tested by far western analysis for their ability to extract PLRV-aphid protein complexes compared to total protein extraction using an SDS denaturing buffer (far right lane). The top panel shows variation in the color of the resulting protein homogenate from cryo-milled aphid tissue extracted in each buffer. Bottom panel shows vector protein bands binding to WT PLRV as determined by one-dimensional separation of extracts followed by incubation with purified WT PLRV (+) and subsequent detection with an in-house PLRV antibody. Negative controls (-) were made by omitting incubation with WT PLRV on a parallel western and these lanes superimposed onto the image of the PLRV (+) blot (gray lines). (B) Western blot analysis of α-PLRV affinity purification experiments from *M. persicae* aphid protein complexes extracted using the TRIS buffer composition show significant enrichment of the PLRV coat protein (CP) and a truncated form (Truncated) of the structural readthrough protein (RTP) in viruliferous (V) *M. persicae* aphids compared to the undiluted affinity purification input fraction and a negative control affinity purification using non-viruliferous insects. Enrichment of the PLRV structural proteins in an AP from systemically infected potato tissue (Pot) is shown as a positive control. Molecular weights corresponding to the full length form of the RTP (FL RTP) and RTP multimers are also indicated.

**Table 1.** Lysis buffer compositions tested for extraction of PLRV-interacting vector proteins.

| Buffer name<sup>a</sup> | Buffer components<sup>b</sup> |
| --- | --- |
| CHAPS | 1X phosphate buffered saline (pH 7.4), 40 mM CHAPS, 10 mM CaCl<sub>2</sub> |
| HEPES | 50 mM HEPES-KOH (pH 7.4), 110 mM KOAc, 2 mM MgCl<sub>2</sub>, 0.4% TritonX-100 |
| TBT | 50 mM HEPES-KOH (pH 7.4), 200 mM Tris (pH7.5), 110 mM KOAc, 350 mM NaCl, 2 mM MgCl<sub>2</sub>, 0.4% TritonX-100, 0.1% Tween-20 |
| TRIS | 50 mM Tris ( pH 7.5), 150 mM NaCl, 0.4% TritonX-100 |
| SDS | 50 mM Tris (pH 6.8), 2.5% Sodium dodecyl sulfate, 10% glycerol |

<sup>a</sup>Names given to protein extraction buffers tested in Fig. 1.
<sup>b</sup>All buffers were supplemented with 0.5 mM phenylmethylsulfonyl and a 1:100 dilution of Halt™ EDTA-free protease inhibitor cocktail. Abbreviations for chemicals are described in materials and methods.

Similar to problems we encountered detecting virus in total protein extracts of systemically infected potato tissue (42), detection of PLRV in viruliferous aphid tissue extracts by western blot analysis using an antibody (α-PLRV) specific for the PLRV structural proteins (40) was too faint and inconsistent to assess extraction efficiency (data not shown). Therefore, we used far-western analysis to gauge the number of vector proteins able to bind with gradient purified PLRV virion in each buffer condition (Fig. 1A, bottom panel). Protein extracts were separated (side-by-side) by one-dimensional SDS-PAGE and transferred to a nitrocellulose membrane and incubated with gradient purified PLRV. Aphid protein bands interacting with virus were subsequently detected with an α-PLRV antibody (Fig. 1A, bottom panel, PLRV+ lanes). A parallel nitrocellulose blot of the same volume of aphid homogenate incubated with only primary and secondary antibodies served as a negative control (Fig. 1A, bottom panel, PLRV-lanes). Using this technique, we identified bands where PLRV bound to similar-sized vector proteins across several extraction conditions and also instances where PLRV bound proteins that differed in molecular weight depending on buffer composition (Fig. 1A, bottom panel). The negative control blot incubated without virus shows that the majority of virus-interacting bands are not the result of cross-reactivity with antibody. Surprisingly, the HEPES-based buffer previously used for plants resulted in zero PLRV-interacting vector protein bands detected above negative control background levels (Fig. 1A, bottom panel, HEPES). Very few PLRV-interacting vector protein bands were detected in the TBT extracted homogenate, most below 37 kilodaltons (kDa) in size. Both the CHAPS-based and Tris-based AP buffers resulted in the extraction of numerous vector protein bands interacting with PLRV virion and/or structural proteins. The Tris-based buffer had a similar protein banding profile as the SDS homogenate positive control and additional PLRV-interacting proteins detected above ∼60 kDa that were not observed in the CHAPS buffer lane.

The Tris-based buffer was chosen for protein extraction in our final affinity purification workflow since it resulted in the greatest number of virus-interacting vector proteins across a wide range of molecular weights. Extraction using this buffer resulted in the faint detection of the PLRV structural proteins, the coat protein (CP, ∼23 kDa) and full-length readthrough protein (RTP, ∼80 kDa) by western blot, in the undiluted homogenate of viruliferous aphids compared to non-viruliferous aphids (Fig. 1B). Affinity purification conducted using α-PLRV conjugated magnetic beads (42) resulted in significant enrichment of both the PLRV CP and a truncated form of the RTP (∼50 kDa) in the AP eluate from viruliferous aphids compared to the undiluted input fraction as shown by western analysis (Fig. 1B). Affinity purification of PLRV from aphids showed a lower level and less diversity of viral structural protein isoforms than affinity purification of PLRV from systemically infected potato homogenate extracted in the HEPES-based buffer (Fig. 1B, Pot). This is indicated by the increased intensity of protein bands corresponding to the PLRV CP and the detection of several different forms of the RTP, including the full-length monomeric form (∼80 kDa), higher molecular weight RTP multimers (>80 kDa), and several RTP truncations (∼50-80 kDa) in the potato affinity purification eluate.

### Assessment of batch effects across independent affinity purification mass spectrometry experiments

The AP-MS experiment was conducted in three independent trials. Each trial consisted of three biological replicate α-PLRV affinity purifications from viruliferous (V) aphids and non-viruliferous (NV) aphids, which served as negative controls for non-specific interaction of aphid proteins with beads and/or antibody. The first two experiments are designated by the month in which the AP batch was conducted (April or January). For the third trial, phosphatase inhibitor was added to the lysis buffer. This experiment is designated as phosphatase inhibitor (PhoIn). To determine the extent of batch effects across our independent experiments and identify deviations in data quality that may affect downstream interaction analyses, we compared the levels of three traditionally used AP-MS quality control metrics (44) to assess technical variability between AP samples (Fig. 2): 1. Abundance of IgG peptides in the samples, 2. MS1 peak area analysis of peptides derived from the bait, in this case CP and RTD, and 3. The total number of proteins identified per affinity purification.

**FIG 2.**
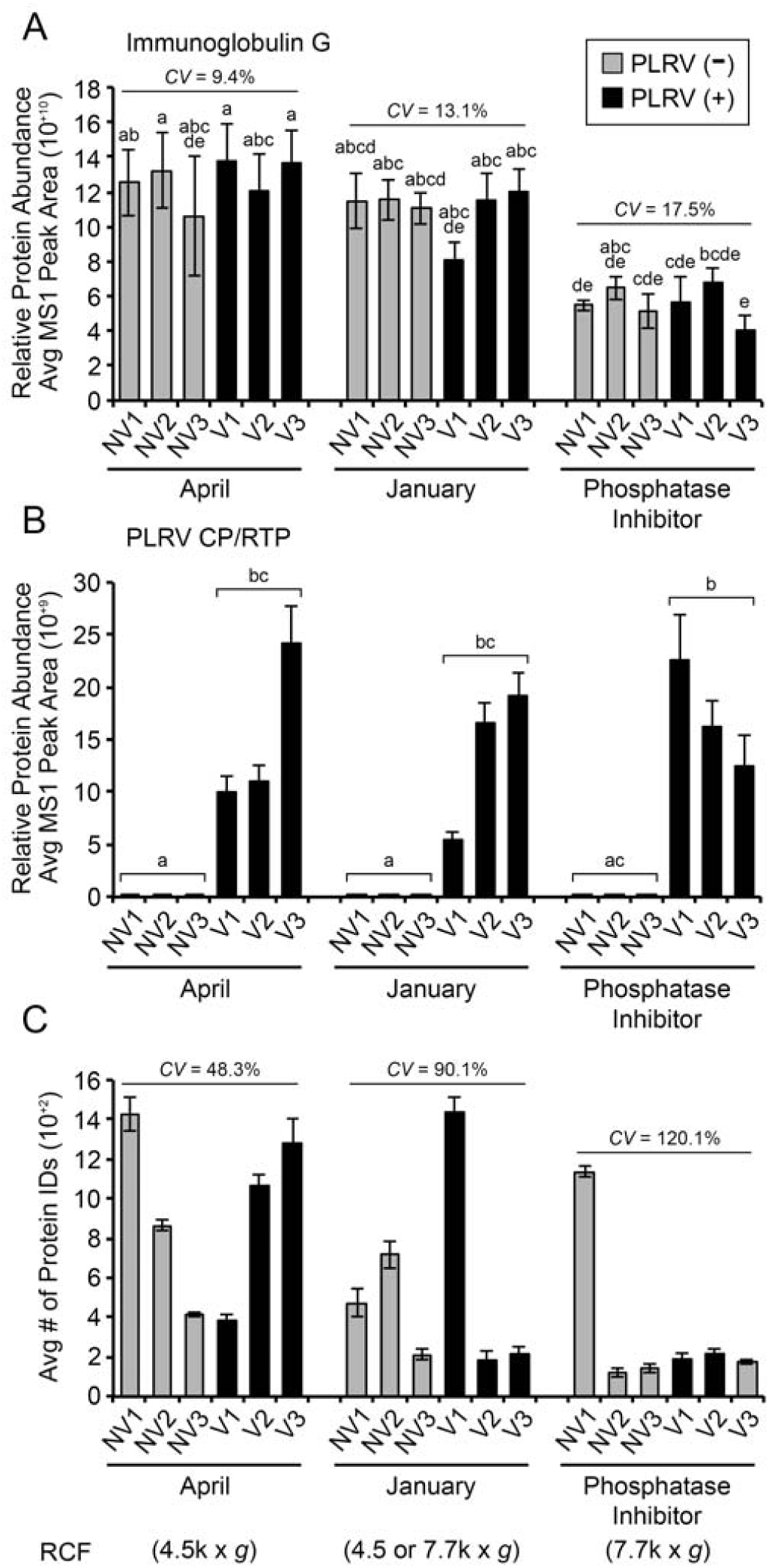
Assessment of variability in α-PLRV affinity purifications from aphids using affinity purification mass spectrometry quality control metrics. (A-B) Bar graphs show the average relative protein abundance of (A) Immunoglobulin G (IgG) and (B) the PLRV structural proteins (CP/RTP) quantified from integration of MS1 (precursor ion) peak areas (unit-less) for protein specific peptides detected in α-PLRV APs from non-viruliferous (NV, gray bars) and viruliferous (V, black bars) aphid pools (*n* =3 analytical replicates per biological replicate) across the three independent datasets (April, January and Phosphatase Inhibitor added). The number of peptides used for quantification of protein abundance are: IgG = 9 and CP/RTP = 7 (Table S2). (C) Bar graph shows the average number of total proteins identified in each biological replicate sample (*n* = 3 analytical replicates) by MS. Centrifugal speed (RCF) of homogenate clarification step is given. For each panel, error bars represent ± one standard error. Lower case letters represent significant differences (*P* < 0.05) calculated by (A) ANOVA and (B) Kruskal-Wallis with Tukey-HSD and Conover (Holm adjustment) post hoc tests, respectively. Lines represent percent coefficient of variance (CV).

**Table 2.**
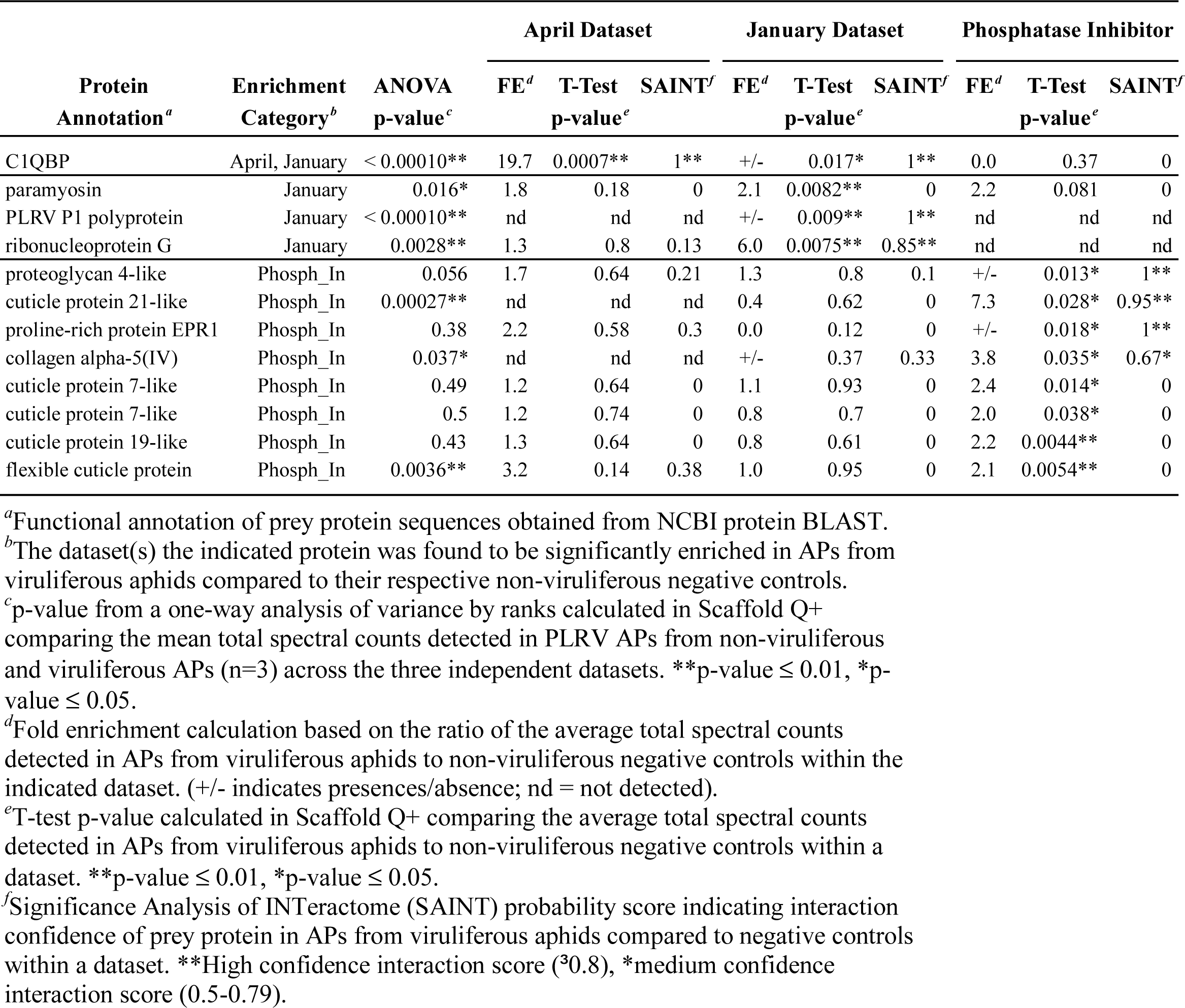
Viral and vector proteins exhibiting high probability of interaction with PLRV in APs from viruliferous aphids using spectral counting.

To assess consistency in the amount of magnetic beads and/or efficiency of antibody conjugation used in each replicate, the relative protein abundance of immunoglobulin G (IgG) was quantified using MS1 peak area integration (Fig. 2A). Within a trial, percent coefficient of variation (CV) of IgG was fairly low (April = 9.4% CV; January = 13.1%; Phosphatase Inhibitor = 17.5%), indicating that, the level of beads/antibody were consistent across all viruliferous (V) and non-viruliferous (NV) biological replicates within an experiment. Levels of IgG in the phosphatase inhibitor dataset were ∼ 2-fold lower as compared to the other datasets.

MS1 peak area integration of peptides specific to the CP and RTD domains of PLRV show that levels of PLRV enrichment in α-PLRV APs from viruliferous aphids were significantly higher compared to negative control APs from non-viruliferous aphids, as expected (Fig. 2B). However, relative variability of PLRV enrichment was high across individual α-PLRV APs from viruliferous aphids within a dataset with %CVs = 52.5 (April), 53.1 (January) and 28.9 (Phosphatase Inhibitor). Interestingly, despite exhibiting a 2-fold decrease in levels of IgG, levels of PLRV in viruliferous biological replicates in the PhoIn dataset had comparable levels of PLRV enrichment, indicating that the lower amounts of beads and/or antibody in these samples did not affect the efficiency of PLRV capture during affinity purification.

Finally, we looked at the average total number of proteins identified per biological replicate (Fig. 2C). Percent CV calculations show a high degree of relative variability across biological replicates within each dataset: 48.3% (April), 90.1% (January), and 120.1% (PhoIn). Differences in the number of proteins identified between the different datasets seem to correlate with the centrifugal speed at which the aphid homogenate was clarified after extraction. The April dataset, where aphid homogenates were cleared at the lowest centrifugation speed, had the highest number of protein identifications across replicates. The PhoIn dataset, in which biological replicates were cleared at the highest centrifugation speed, had the lowest number of protein identifications with the exception of the non-viruliferous, biological replicate 1 (Fig. 2C, Phosphatase Inhibitor, NV1). The January dataset had one biological replicate of viruliferous aphids (Fig. 3C, January, V1), which had the highest number of proteins identified, but also had a very low enrichment of PLRV (Fig. 2B, January, V1). In general, these results show that more proteins were identified when centrifugal speeds were slower and/or when bait capture was low, suggesting that variability in total protein identification may have been a result of increased non-specific binding to antibody and/or beads.

**FIG 3.**
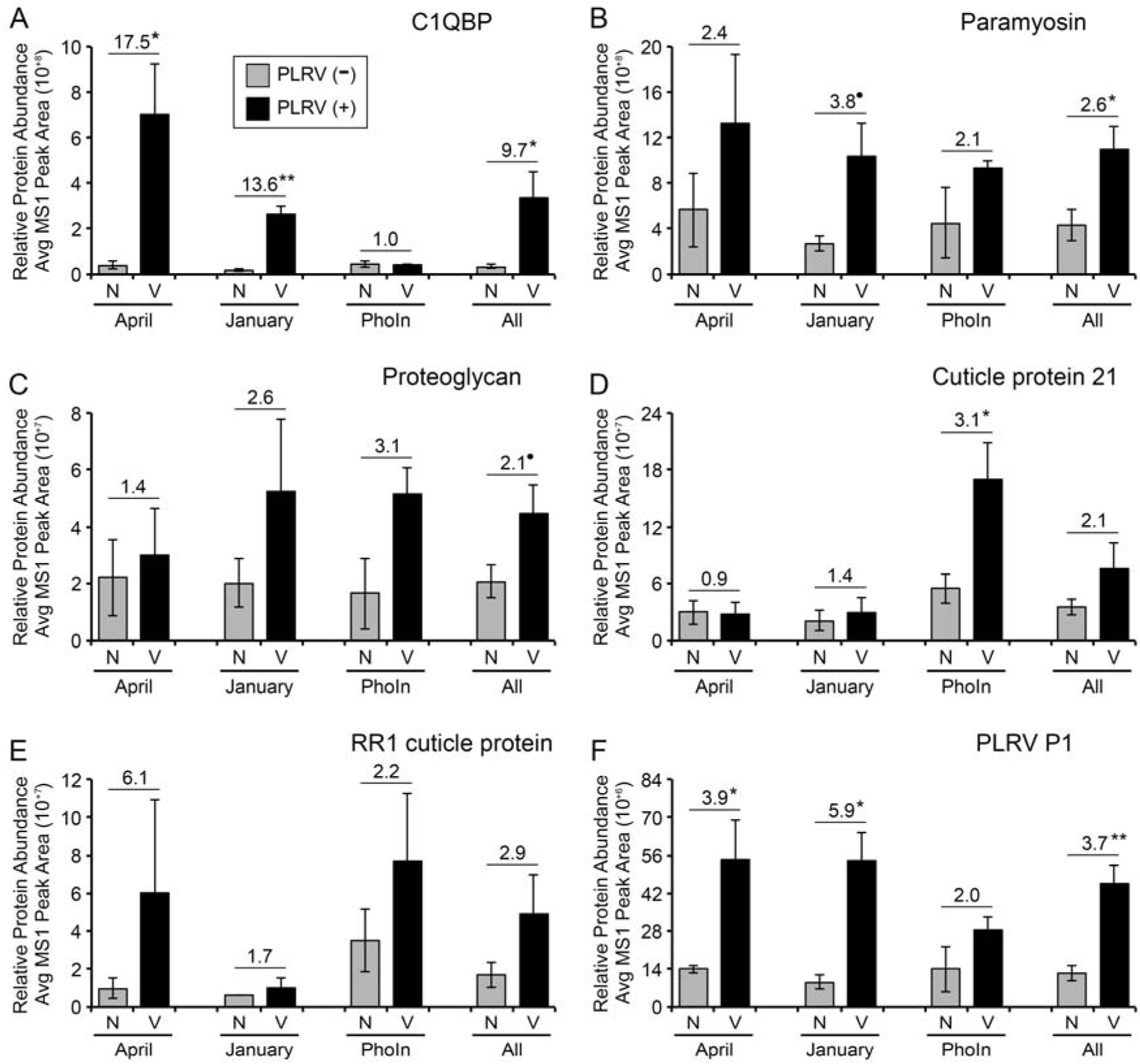
Label-free quantification of vector and viral protein enrichment in α-PLRV affinity purifications from viruliferous *M. persicae* using MS1 peak integration. Bar graphs show relative protein abundance measured by integration of MS1 (precursor ion) peak areas (unit-less) of protein specific peptides corresponding to a selected group of (A-E) vector and (F) viral proteins found to have significantly enriched total spectral counts in α-PLRV affinity purification biological replicates from viruliferous aphids (Table 1 and 2). The number of peptides used for MS1 quantification are (A) C1QBP = 6, (B) Paramyosin = 21, (C) Proteoglycan-4 like = 4, (D) Cuticle protein 21 = 6, (E) RR1 cuticle protein 5 = 3 and (F) the PLRV P1 polyprotein = 3. The MS1 integration data is shown as an average of *n* = 3 biological replicate α-PLRV APs from non-viruliferous aphids (NV, gray bars) and viruliferous aphids (V, black bars) compared within each of the three independent datasets: April, January and phosphatase inhibitor added (PhoIn). The average protein abundance for all three datasets combined (All, *n* = 9 biological replicates) is also shown. Error bars represent ± one standard error. Lines above bars indicate fold enrichment in α-PLRV affinity purifications from viruliferous aphids compared to affinity purifications from non-viruliferous aphids with statistical significance (• = *P* < 0.06, * = *P* < 0.05, ** = *P* < 0.01) calculated by Student’s *t*-test (normal data) or Welch’s T-test (non-normal data) within an AP dataset.

### Identification of viral and insect proteins *in complex* with PLRV isolated from viruliferous aphids by affinity purification

Due to the variability between datasets, we decided to perform label-free quantification of the peptides from each dataset separately to identify vector proteins that were significantly enriched with virus captured from viruliferous aphids compared to non-viruliferous controls. Initially, we identified a total of 106 vector proteins or protein clusters and one non-structural viral PLRV protein, the P1 polyprotein, whose average total spectral counts (SPC) were enriched ≥ 2-fold or present/absent (+/-) in viruliferous APs in one or more of our three datasets compared to their respective non-viruliferous controls (Table S1). Taking into consideration the high variability observed in bait levels and total protein identification across AP samples, another filter criterion was that the enriched prey protein had to be detected in at least two of the three viruliferous AP biological replicates within a dataset. From this refined list of putatively enriched proteins, the average total SPC of 11 vector proteins and the PLRV P1 polyprotein were found to be significantly enriched in viruliferous APs by one or more statistical analyses: T-test, One-way ANOVA, and/or had a high-confidence probability score of ≥ 0.8 using the AP-MS statistical tool Significance Analysis of INTeractome (SAINT), a program that utilizes negative control AP data to identify non-specific interactions in a semi-supervised manner and computes confidence scores (probability) for putative interactions (45–47). We categorized these as high probability candidate interactions (Table 2). We identified a second category of 23 vector proteins (Table 3) with average total SPCs ≥ 2-fold enriched or +/− in viruliferous APs in only one AP dataset but had medium confidence SAINT interaction scores (0.5 to 0.79).

**Table 3.** Vector proteins exhibiting medium confidence interaction scores with PLRV using spectral counting.

| Protein<br>Annotation<sup>a</sup> | Enrichment<br>Category<sup>b</sup> | April |  | January |  | Phosphatase<br>Inhibitor |  |
| --- | --- | --- | --- | --- | --- | --- | --- |
|  |  | FE<sup>c</sup> | SAINT<sup>d</sup> | FE<sup>c</sup> | SAINT<sup>d</sup> | FE<sup>c</sup> | SAINT<sup>d</sup> |
| cuticle protein 7-like | April | 6.5 | 0.62* | 0.4 | 0 | 2.2 | 0.4 |
| possible chitinase | April | +/- | 0.63* | 0.0 | 0 | 2.1 | 0.32 |
| putative inner membrane peptidase | April | +/- | 0.65* | nd | nd | nd | nd |
| splicing factor U2AF 50 kDa subunit | April | +/- | 0.65* | nd | nd | nd | nd |
| endothelin-converting enzyme | April | +/- | 0.65* | +/- | 0.33 | 0.0 | 0 |
| MARK2-like isoform X1 | April | +/- | 0.65* | nd | nd | 0.0 | 0 |
| coiled-coil domain-containing protein | April | +/- | 0.65* | nd | nd | nd | nd |
| RNA-binding protein cabeza-like | April | 6.0 | 0.61* | 4.0 | 0.33 | 0.0 | 0 |
| cuticle protein 16.5 | January | 0.9 | 0.02 | 3.2 | 0.64* | 2.4 | 0.04 |
| DNA-directed RNA polymerase II | January | 1.4 | 0.05 | +/- | 0.67* | 1.7 | 0 |
| papilin isoform X1 | January | 1.0 | 0.05 | +/- | 0.67* | 0.0 | 0 |
| 60S acidic ribosomal protein | January | 1.2 | 0 | +/- | 0.67* | 0.0 | 0 |
| U4/U6.U5 tri-snRNP-associated protein | January | nd | nd | +/- | 0.66* | 1.5 | 0.22 |
| small nuclear ribonucleoprotein D2-like | January | 1.1 | 0.02 | +/- | 0.66* | nd | nd |
| uncharacterized protein LOC111032353 | January | nd | nd | +/- | 0.65* | 2.7 | 0.31 |
| prostaglandin reductase 1-like | January | 0.8 | 0 | 2.6 | 0.60* | 1.3 | 0 |
| RR1 cuticle protein 5 | Phosph_In | 4.2 | 0.34 | nd | nd | 15.5 | 0.66* |
| HSPG2 | Phosph_In | 0.0 | 0 | 2.4 | 0.5 | 5.0 | 0.62* |
| proline-rich protein 36-like | Phosph_In | 1.9 | 0.3 | 0.0 | 0 | 6.8 | 0.73* |
| uncharacterized protein LOC111041433 | Phosph_In | nd | nd | +/- | 0.33 | +/- | 0.67* |
| tubulointerstitial nephritis antigen | Phosph_In | nd | nd | 1.1 | 0.14 | +/- | 0.65* |
| proline-rich protein EPR1 | Phosph_In | nd | nd | 1.0 | 0.04 | +/- | 0.65* |
| nuclear transcription factor Y subunit | Phosph_In | 1.0 | 0.15 | +/- | 0.33 | +/- | 0.65* |

<sup>27</sup>
<sup>a</sup>Functional annotation of prey protein sequences obtained from NCBI protein BLAST.
<sup>b</sup>The dataset(s) the indicated protein was found to be significantly enriched in APs from viruliferous aphids compared to their respective non-viruliferous negative controls.
<sup>c</sup>Fold enrichment calculation based on the ratio of the average total spectral counts detected in APs from viruliferous aphids to non-viruliferous negative controls (n=3) within the indicated dataset. (+/- indicates presences/absence; nd = not detected).
<sup>d</sup>Significance Analysis of INTeractome (SAINT) probability score indicating interaction confidence of prey protein in APs from viruliferous aphids compared to negative controls within a dataset. \*Medium confidence interaction score (0.5-0.79).

Within the high probability interaction group, C1QBP was the only vector protein identified as significantly enriched in viruliferous APs in multiple datasets: 19.2-fold enrichment in the April dataset and present/absent (+/-) in the January dataset (Table 2 and Table S1). Zero spectral counts were detected in viruliferous APs in the dataset where phosphatase inhibitor was added (Table S1), although some spectral counts were detected in the APs of non-viruliferous aphids within the same dataset. The remaining 10 vector proteins as well as the PLRV P1 polyprotein were found significantly enriched in either the January or PhoIn dataset. Five of these proteins were annotated as cuticular proteins and had average total SPCs that were significantly enriched in the viruliferous APs where phosphatase inhibitor was added to the extraction buffer.

Since spectral-based counting is dependent upon machine selection of peptide signals for MS^2^ fragmentation, it is often biased towards under-sampling low abundant peptides (48). Therefore, we used a second label-free quantification method, MS1 peak integration, to measure the full precursor ion signal intensities (MS^1^) for peptides corresponding to a subset of these putative PLRV-interacting proteins to validate their enrichment trends in viruliferous APs compared to their respective non-viruliferous controls (Fig. 3, Table S2). Significant enrichment of C1QBP was confirmed in the APs from viruliferous aphids in the April and January datasets (Fig. 3A). Levels of C1QBP were low and equal in AP samples where phosphatase inhibitor was added. Two vector proteins, paramyosin and a proteoglycan 4-like protein, both predicted to be high probability candidate interactions in the January or PhoIn spectral counting datasets, respectively (Table 2), did exhibit an average trend of MS1-based enrichment (1.4 to 3.8-fold) in viruliferous APs compared to their respective non-viruliferous controls in all three datasets (Fig. 3B-C). Average co-enrichment for paramyosin was statistically significant (*P* < 0.05, Student’s *t*-test) when biological replicates from all three datasets were averaged together (Fig. 3B, All) and *P* = 0.059 for the January dataset (Fig. 3B, January, Student *t*-test). Similarly, when all biological replicates were analyzed together, the significance of enrichment for the proteoglycan-like protein was *P* = 0.058 (Fig. 3C, All, Student *t*-test). Comparable to what was observed by spectral-counting, *M. persicae* cuticle protein 21 was significantly enriched (*P* < 0.05, Student *t*-test) only in the dataset where phosphatase inhibitor was added (Fig. 3D, PhoIn). However, an RR1-like cuticle protein from our medium-confidence interaction group, even though it exhibited a trend of enrichment in viruliferous APs in all three datasets, as the differences were not significant (Fig. 3E). Lastly, significant enrichment of the PLRV P1 polyprotein was confirmed by MS1 peak integration in viruliferous APs in both the April and January dataset (Fig. 3F), even though spectra-based counting indicated no sampling of P1 in the April APs (Table 2). This was most likely due to interference of a co-eluting peptide within the same retention time window (data not shown) as indicated by background signal detected in non-viruliferous AP controls even though aphids were fed on plants that did not contain virus (Fig. 3F, NV).

### Validation of direct interaction between luteovirid structural proteins and vector proteins identified by AP-MS using yeast-two-hybrid

In a parallel experiment, a yeast-two-hybrid screen of a cDNA library constructed from whole-body mRNA extracts of the green bug aphid *S. graminum,* a competent vector of the yellow dwarf viruses, confirmed direct interaction of luteovirid structural proteins with orthologues of three *M. persicae* proteins identified in our AP-MS experiment. *S. graminum* proteins translated from vector expressed sequence tags (ESTs) fused to the activating domain (AD) of the yeast transcription factor GAL4, were screened for interaction with GAL4 DNA binding domain (BD) protein fusions of the CP or readthrough domain (RTD) from the luteovirids *Barley yellow dwarf virus-PAV* (BYDV-PAV, genus *Luteovirus*), *Cereal yellow dwarf virus-RPV* (CYDV-RPV, genus *Polerovirus*), and PLRV, which *S. graminum* does not transmit. From this initial screen and subsequent co-transformation experiments, a partial sequence of SgC1QBP (∼98% identity to C1QBP from *R. madis,* XP_026823488.1) was found to directly interact with the CP of BYDV-PAV, CYDV-RPV, and PLRV as indicated by blue coloring and growth on quadruple dropout media supplemented with X-α-gal. Negative controls showed that AD-SgC1QBP did not interact with BD protein fusions of murine p53 and lamin C (Fig. 4A).

**FIG 4.**
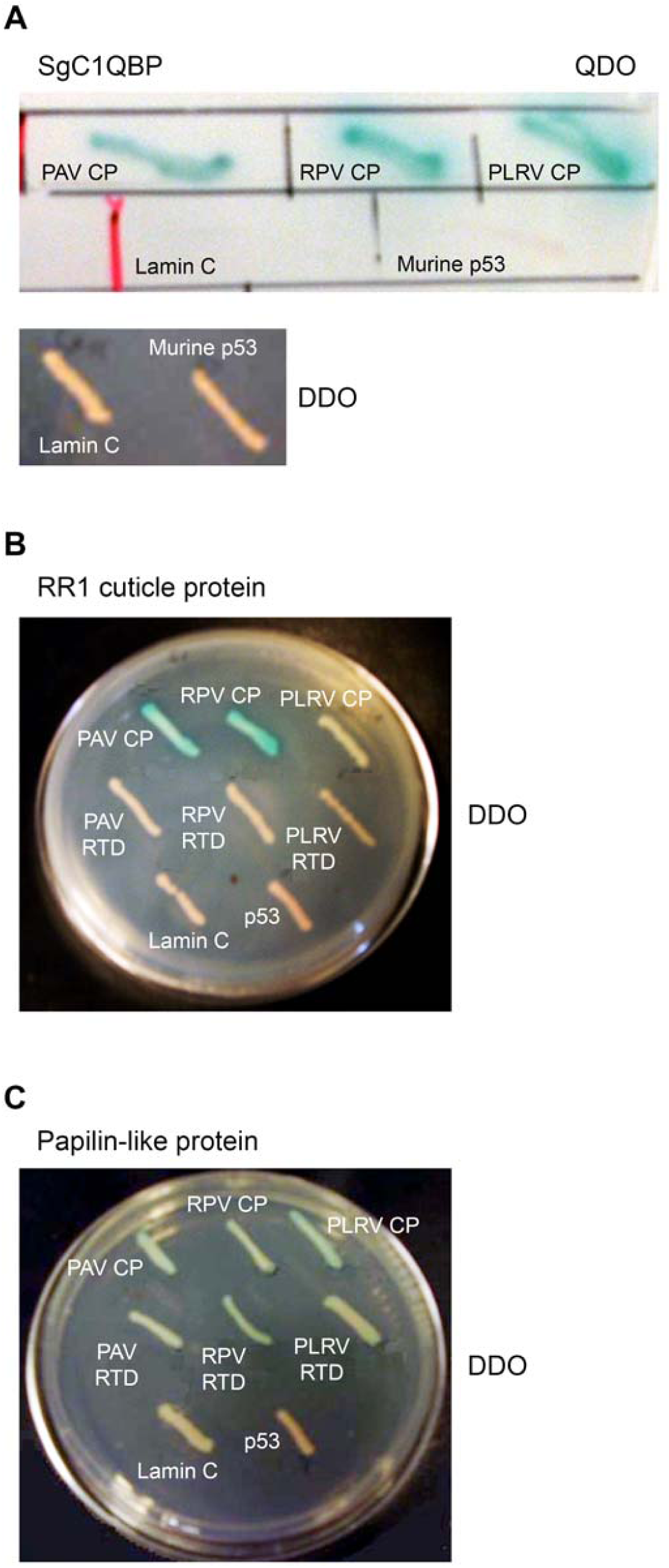
Identification of *Schizaphis graminum* proteins directly binding to luteovirid structural proteins using yeast-two-hybrid (Y2H) assay. Interactions between (A) C1QBP, (B) RR1-type cuticle protein and (C) papilin-like protein from *S.s graminum* (Sg) with the coat protein (CP) or readthrough domain (RTD) of the luteovirids BYDV-PAV (PAV), CYDV-RPV (RPV), and PLRV. Vector and viral proteins were co-expressed as GAD-T7-Rec and DNA-BD fusions, respectively. Co-transformed cells were grown on yeast quadruple-dropout (QDO) medium (SD/-Ade/-His/-Leu/-Trp) and/or double-dropout (DDO, SD/-Leu/-Trp) supplemented with X-α-Gal. Co-transformation of GAD-T7-Rec-vector protein constructs with lamin C or murine p53 fused to DNA-BD were used as negative controls.

Two other *S. graminum* proteins were also identified from this Y2H screen that overlapped in function with vector proteins identified as enriched in our AP-MS experiment with PLRV: a RR1-type cuticle protein and papilin isoform X1 (Table 3). A 280 nucleotide (nt) EST coding for a cuticle protein belonging to the RR1 group and a 560 nt EST sequence showing similarity to an ectodermal papilin-like protein (XP_015371026.1), on the amino acid level were identified during the screening of the full body *S. graminum* cDNA library with the CP of CYDV-RPV. The *S. graminum* RR1 cuticular protein was found to directly interact with the CP of CYDV-RPV and BYDV-PAV indicated by the blue-coloring of co-transformants grown on double dropout media, but not the CP of PLRV, nor the RTD sequences of any of the viruses (Fig. 4B). The *S. graminum* papilin-like protein was found to weakly interact (Fig. 4C, fainter blue coloring) with both virus structural protein domains, CP and RTD, from BYDV-PAV, CYDV-RPV, and PLRV. Again, no interaction was detected with the negative control p53 and lamin C proteins with either of these vector proteins, indicating the interactions to be specific. Interestingly, homologs of both these proteins were assigned medium-confidence interaction SAINT scores (0.66 and 0.67, respectively) in only one dataset of our *M. persicae* AP-MS experiments (Table 3). Furthermore, label-free quantification of relative protein abundance by MS1 peak integration showed that, although the RR1 cuticle protein homolog had a trend of enrichment in all AP-MS datasets, high variability in protein levels across APs most likely reduced our ability to detect statistical significance, a scenario that often occurs when protein interactions are low abundant, weak and/or transient (49). Collectively, our data highlight the need for complementary experimental approaches to validate these types of protein-protein interactions that may fall short of accepted significance thresholds in challenging AP-MS experiments where bait levels are low and variability (technical and/or biological) is high.

### Measuring changes in the relative abundance of PLRV-interacting aphid proteins after feeding on an infected host plant

We used a shotgun proteomics approach to assess whether the abundance of aphid proteins predicted to complex with PLRV by AP-MS (Tables 2 and 3) changed their abundance during virus acquisition. Within this shotgun dataset, peptides from six of our candidate PLRV-interacting *M. persicae* proteins were detected: C1QBP, paramyosin, collagen alpha-5 chain, cuticle protein 21, RR1 cuticle protein and basement membrane-specific heparan sulfate proteoglycan core protein (HSPG2).

Quantification by spectral-based counting showed that the average relative abundance of five of these proteins, including our highest confidence PLRV-interacting protein, C1QBP, were unchanged between viruliferous and non-viruliferous aphids (Fig. 5), indicating that their increase in abundance in α-PLRV APs from viruliferous aphids was not a consequence of higher expression of a protein non-specifically interacting with the antibody-coated magnetic beads. Conversely, one of the vector proteins predicted to form a high-confidence interaction with PLRV, collagen alpha-5 chain, which exhibited 3.8-fold enrichment (average total SPC) in viruliferous APs where phosphatase inhibitor was added (Table 2), also shows a 2-fold increase in expression in viruliferous insects (Fig 5C). With our current data, it is unclear whether the enrichment of collagen alpha-5 chain that was detected in viruliferous APs is solely due to a greater abundance of this protein in viruliferous insects or if it is a true interaction. Regardless, these results show that acquisition of PLRV by *M. persicae* leads to the differential expression of this protein.

**FIG 5.**
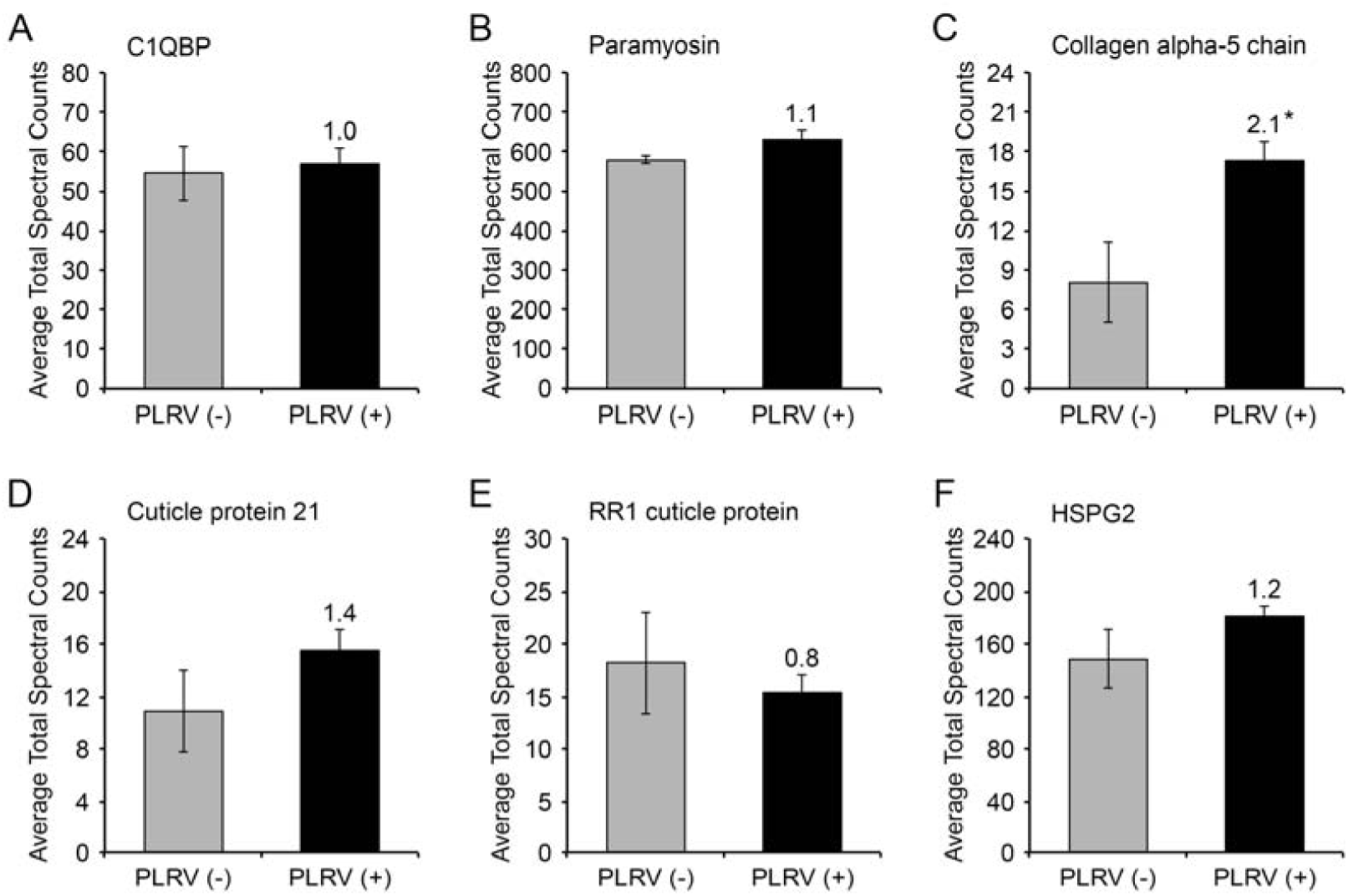
PLRV interacting proteins in viruliferous compared to non-viruliferous aphids do not change expression in aphids upon virus acquisition. Graphed are the average total spectral counts (SPC) of (A) C1QBP, (B) Paramyosin, (C) Collagen alpha-5 chain, (D) Cuticle protein 21, (E) RR1 cuticle protein and (F) HSPG2 in viruliferous (PLRV +, black bars) or non-viruliferous (PLRV -, gray bars) *M. persicae* aphids (*n* = 3-4 pools of aphids). Values above black bars indicate fold enrichment in viruliferous aphids compared to non-viruliferous controls and significantly different groups via Student’s t-test (* = *P* < 0.05) and error bars represent ± one standard error.

Collectively, the proteomics and yeast-two hybrid experiments are hypothesis-generating tools. The most statistically enriched vector protein in our AP-MS experiments is MpC1QBP, which also directly binds to the CP of luteovirids. We decided to test the hypothesis that MpC1QBP was involved in PLRV transmission by *M. persicae* by performing additional experiments to characterize this protein and test whether it regulates virus transmission.

### C1QBP from *M. persicae* shares sequence similarity and conserved truncation sites with C1QBP from humans and other insects

C1QBP is highly conserved among aphid species with most sharing >90% amino acid sequence identity with the C1QBP sequence from *M. persicae*, but the protein is less conserved among other distantly related hemipterans such as the Asian citrus psyllid and planthopper species (Fig. 6). Human C1QBP (28% identity /43% similarity to MpC1QBP) is a mitochondrial matrix protein (p32) involved in a broad array of pathways including immunity and cancer progression and has been shown to localize to multiple cellular compartments (52–55). The yeast orthologue, MAM33, is a molecular chaperone that functions in the assembly of mitoribosomal proteins, which regulate the translation of key proteins involved in oxidative phosphorylation (56).

**FIG 6.**
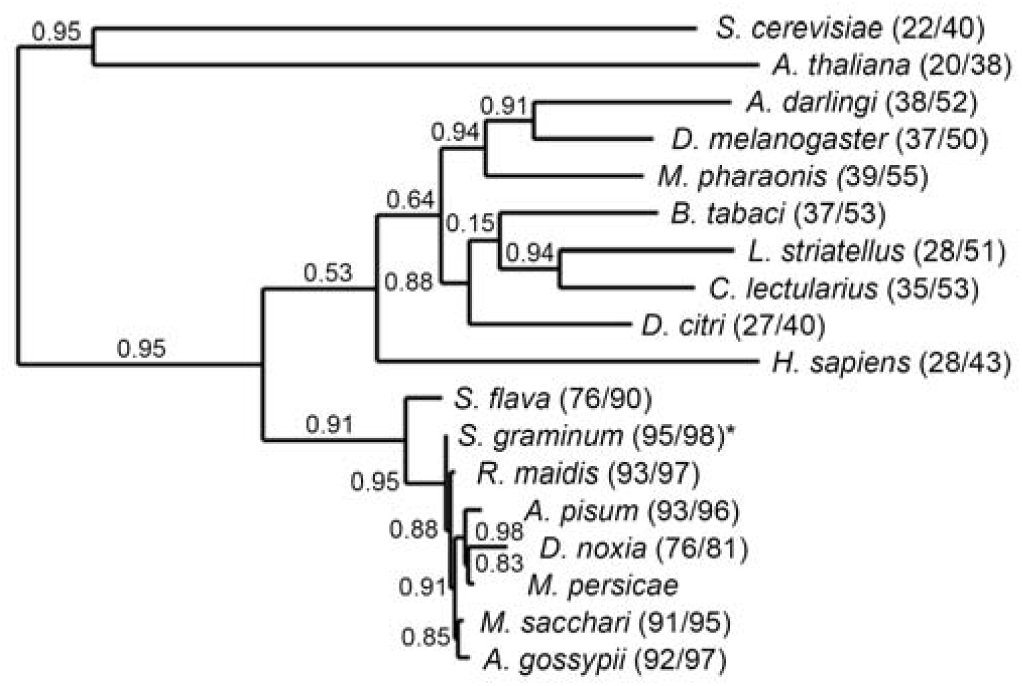
Phylogenetic analysis of C1QBP orthologous protein sequences using maximum likelihood indicates C1QBP from insects is more closely related to C1QBP from humans than yeast or plants. The analysis included 18 protein sequences representing aphids, insects and other diverse model organisms. Branch points and bootstrap values were obtained from 100 iterations using the PhyML 3.1/3.0 aLRT algorithm and TreeDyn for tree drawing. Values in parentheses represent the percent identity/percent similarity in a pairwise sequence comparison with the *M. persicae* C1QBP sequence using BLAST Global Align. Accession numbers for all sequences used in this analysis are listed in Materials and Methods.

Global alignment with MpC1QBP of representative orthologous sequences across kingdoms shows high conservation of residues in the C-terminus of the protein (amino acids 187-243) but less so in the N-terminal region of these proteins (Fig. 7A). In humans, the first 73 amino acids of the C1QBP N-terminus codes for a mitochondrial targeting peptide that is cleaved after import into the mitochondrial matrix (57). The two residues immediately following the known truncation position in human C1QBP (Fig. 7A, red arrowhead) are highly conserved between human and insect sequences (*D. melanogaster, A. pisum,* and *M. persicae*), with all four species having a histidine residue immediately downstream of the truncation site followed by a serine in aphids or the biochemically similar residue, threonine, in *D. melanogaster* and humans. These residues are not conserved at this position in C1QBP orthologues from yeast and plants. We did not identify peptides corresponding to the MpC1QBP N-terminus upstream of this predicted truncation site in any of our APs from viruliferous aphids (Fig. 7B), even though tryptic peptides are predicted to be produced from this region. The first tryptic peptide identified in our data, K.ELQQFLDNEIKSEEQTSDK.S, is located six residues downstream of the predicted truncation site (Fig. 6B, red box). Furthermore, the *S. graminum* Y2H screen also only identified an EST that produced a truncated form of C1QBP (Fig. 6, blue arrowhead). These data suggest that PLRV most likely interacts exclusively with the truncated form of C1QBP *in vivo*.

**FIG 7.**
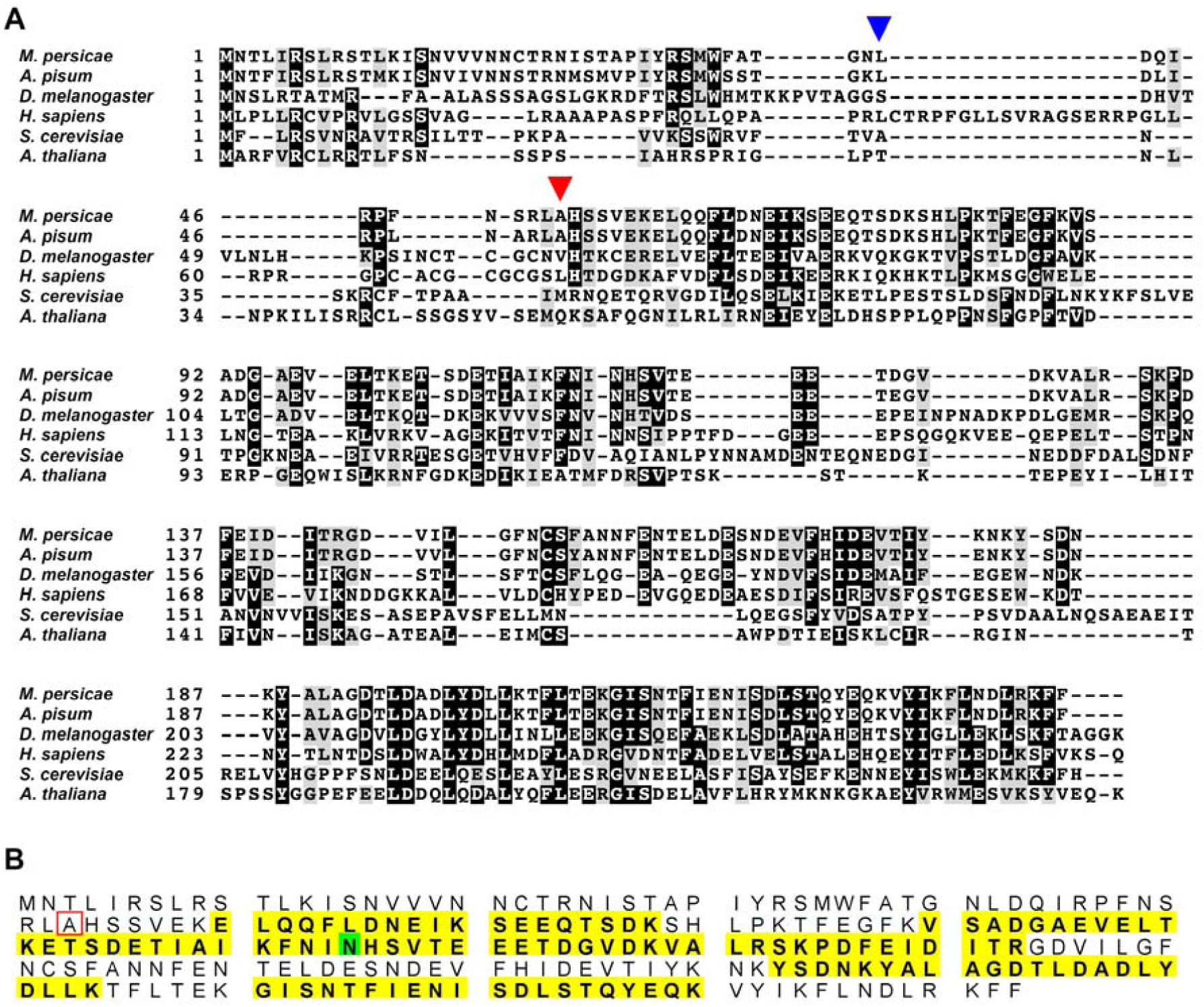
Multiple sequence alignment of C1QBP proteins from diverse organisms shows conservation of an N-terminal truncation site and the C-terminal region of the protein. (A) Selected orthologous C1QBP protein sequences from aphids (*M. persicae* and *A. pisum*), fruit flies (*D. molenogaster),* humans (*H. sapiens*), yeast (*S. cerevisiae*) and plants (*A. thaliana*) were aligned using Clustal Omega Multiple Sequence alignment and visualized using BoxShade. Black boxes indicate identical residues whereas gray boxes highlight residues that are similar. The red arrowhead highlights the known site of N-terminal truncation of the human form of C1QBP. The blue arrowhead indicates the residue corresponding to the start of the truncated *S. graminum* C1QBP protein that was identified interacting with the luteovirid BYDV-RPV by yeast-two-hybrid screening. (B) Visual representation of the peptide coverage for the *M. persicae* C1QBP protein found to be significantly co-enriched with PLRV. Yellow blocks indicate the tryptic peptides detected by nanoflow LC-MS/MS after analysis in Scaffold. The green block highlights a deamidated asparagine residue. The alanine residue corresponding to the known site of N-terminal truncation in human C1QBP is highlighted by a red box.

### C1QBP and PLRV localize to distinct but overlapping subcellular compartments in *M. persicae* midgut cells

Whole-mount immunostaining of guts dissected from viruliferous aphids with an antibody raised against the full-length version of human C1QBP (α-HsC1QBP) and α-PLRV showed MpC1QBP signal could be detected by laser scanning confocal microscopy throughout the alimentary canal (anterior midgut, posterior midgut, and hindgut) and overlapped with α-PLRV signal in the midgut (Fig. 8A). The subcellular localization pattern of α-HsC1QBP signal was mainly observed as diffuse puncta within the cytoplasm of midgut cells, which partially co-localized with α-PLRV signal in some areas (Fig. 9B-C, white arrows). In areas where puncta do not co-localize, fluorescence from α-HsC1QBP and α-PLRV were often adjacent to one another suggesting that subcellular compartments containing C1QBP may be coming into contact with those that contain PLRV (Fig. 8C). Intriguingly, α-HsC1QBP labeled puncta could also be observed aligned along the cell periphery, again partially co-localizing or adjacent to α-PLRV labeled puncta (Fig. 8D-E, white arrowheads). Fluorescence was not observed in viruliferous guts stained with secondary antibodies only (Fig. 8F). Nor was fluorescence signal from α-PLRV detected in non-viruliferous guts imaged with the same parameters (Fig. 8G) indicating that α-PLRV signal was specific. Furthermore, the localization pattern of α-HsC1QBP in non-viruliferous guts was similar to that found in viruliferous aphid cells (Fig. 8G). Our data support the hypothesis that C1QBP and PLRV are in proximity to directly interact in aphid gut epithelial cells, potentially along the cell periphery, plasma membrane and in the cytoplasm.

**FIG 8.**
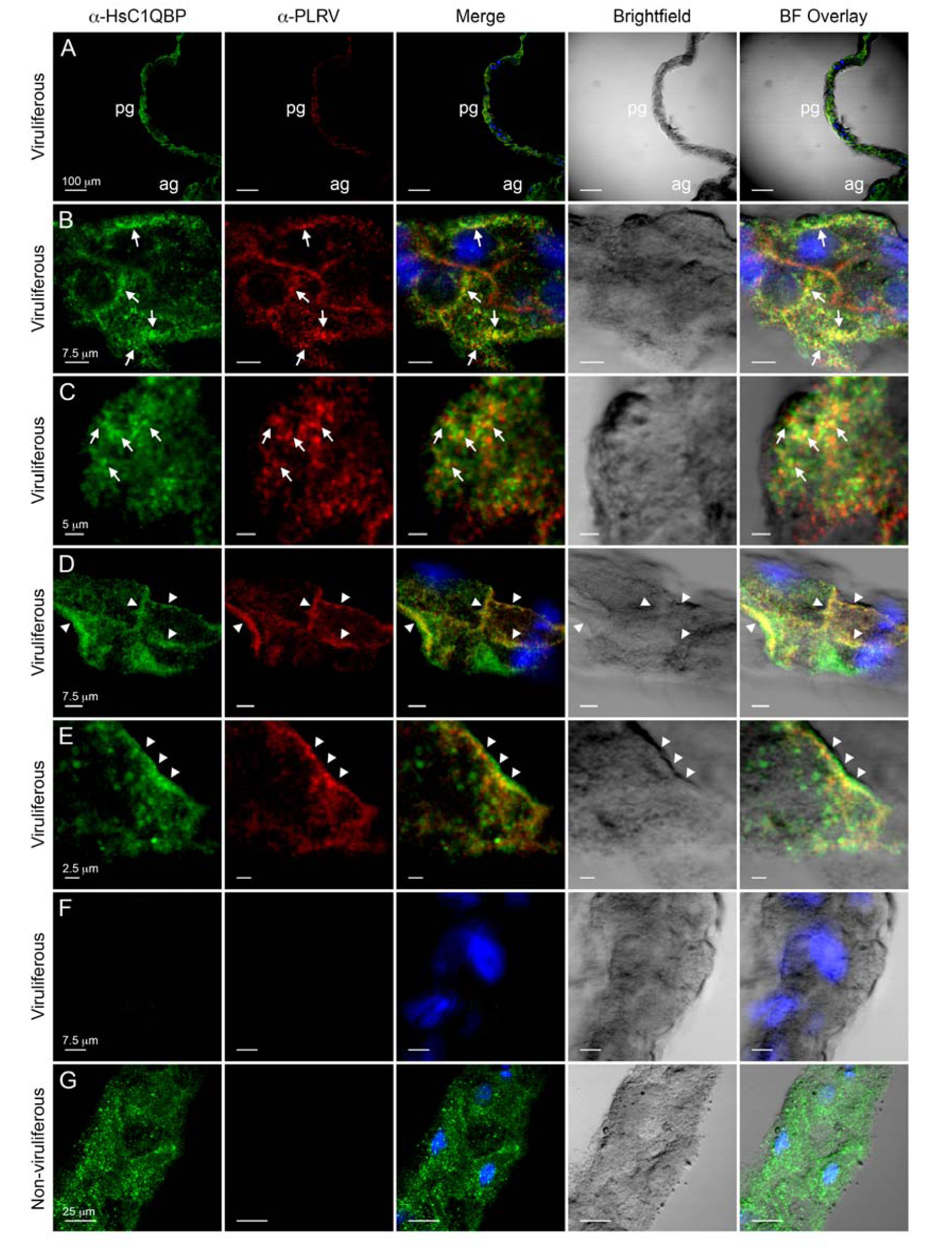
Immunolocalization of C1QBP and PLRV in *M. persicae* gut epithelial cells. Panels show representative single-plane confocal micrographs of guts dissected from viruliferous or non-viruliferous aphids that were immunolabeled with both α-HsC1QBP (Cy2, green) and α-PLRV (Cy3, red). The overlap of the fluorescent signals from α-HsC1QBP and α-PLRV is shown in the column labeled Merge with co-localization appearing in yellow. Nuclei stained with DAPI appear blue. Brightfield images and the Brightfield (BF) overlay are also shown. (A) 10x magnification of the anterior midgut (ag) and posterior midgut (pg) of a single viruliferous gut shows localization of α-HsC1QBP throughout the entire midgut while the fluorescence indicating α-PLRV is strongly detected in the posterior midgut. (B-E) Within individual posterior midgut cells from different guts, points of co-localization of α-HsC1QBP and α-PLRV could be observed as diffuse puncta (B-C) always within the cytoplasm (white arrows) or (D-E) sometimes at the cell periphery (white arrowheads). (F) Incubation of a viruliferous gut with only Cy2-conjugated and Cy3-conjugated secondary antibodies and (G) non-viruliferous gut with both α-HsC1QBP and α-PLRV represent our negative controls. The Cy2 signal in panels (C) and (E) was imaged using a hybrid (HyD) detector and a PMT detector in all other images. Scale bars equal the length indicated.

**FIG 9.**
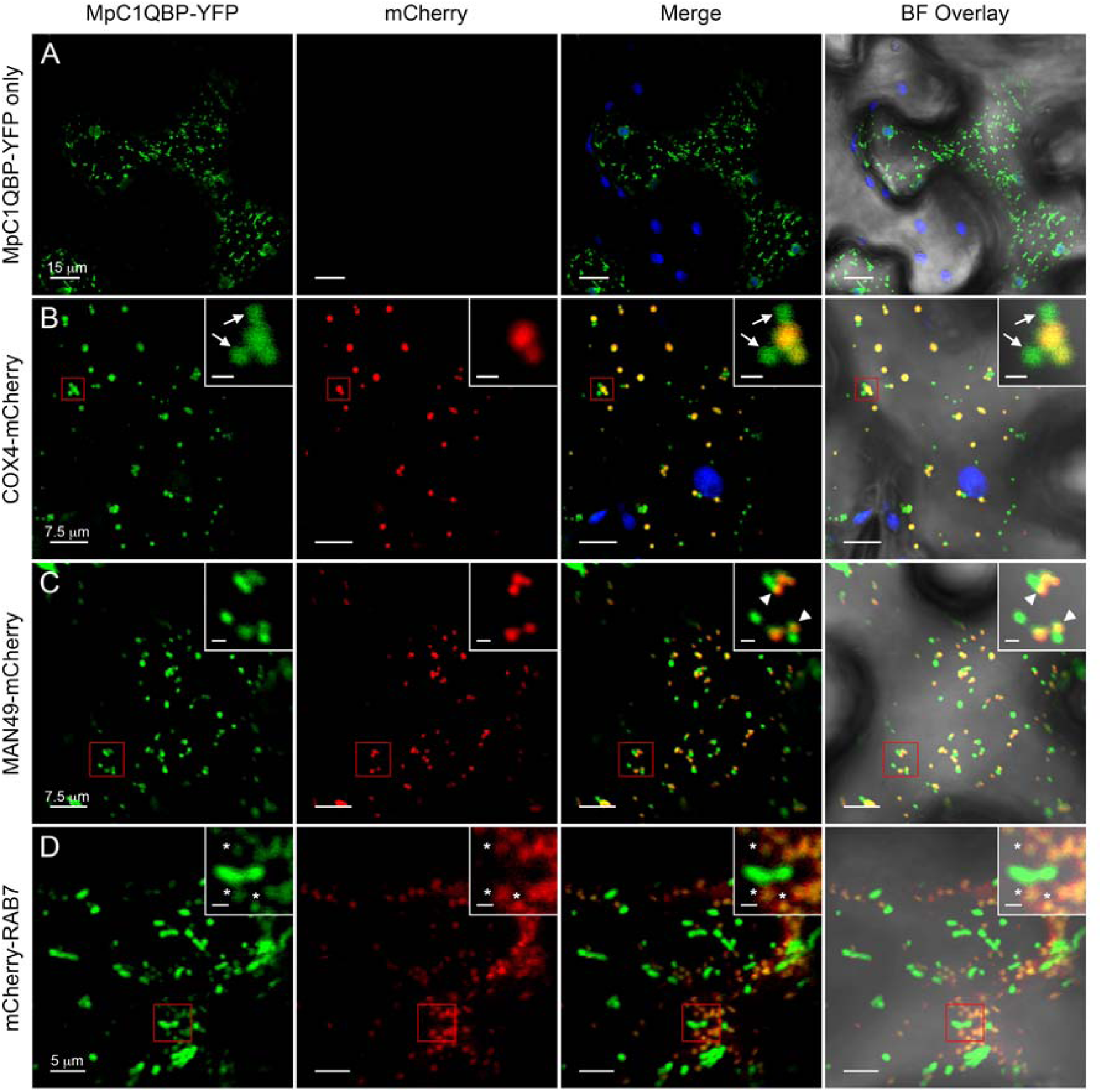
Heterologous expression of *M. persicae* C1QBP in plants shows localization to multiple, motile organelles similar to localization in human cells. Panels show representative single-plane confocal micrographs of live *N. benthamiana* epidermal cells constitutively expressing (A) MpC1QBP-YFP alone or with the organelle markers (B) COX4-mCherry (mitochondria), (C) MAN49-mCherry (*cis*-Golgi) or (D) mCherry-RAB7 (transitory late endosomes). The fluorescence from MpC1QBP-YFP and mCherry are false colored green and red, respectively. The overlap of the fluorescent signals from YFP and mCherry is shown in the column labeled Merge with co-localization of the indicated fusion proteins appearing in yellow. The brightfield overlay (BF Overlay) is also shown. Chloroplast autofluorescence is falsely colored blue. The inset in each panel is a magnified view of the area highlighted by the red box. White arrows indicate globular, MpC1QBP-YFP fluorescence that was observed fusing with mitochondrial-localized MpC1QBP-YFP. White arrowheads mark the sites of partial co-localization of MpC1QBP-YFP with the MAN49-mCherry labeled *cis*-Golgi. White asterisks highlight mark some sites of MpC1QBP-YFP and mCherry-RAB7 co-localization. Scale bars within the main panels show the length indicated while inset scale bars equal 1 μm.

### Localization of MpC1QBP to motile cellular compartments is conserved

In addition to mitochondria, human C1QBP has also been shown to localize to several other subcellular compartments including endosomes and the plasma membrane (58–62), two compartments within cells that PLRV uses to transverse the aphid gut. Such detailed cellular studies are limited in aphids where ectopic expression of markers is currently not possible. Therefore, to assess conservation of subcellular localization that could be correlated with the localization patterns observed in aphid guts, we ectopically co-expressed an MpC1QBP-YFP fusion protein in the model plant *Nicotiana benthamiana* with or without several previously described fluorescently tagged organelle markers: COX4-mCherry (mitochondria), MAN49-mCherry (Golgi), and mCherry-RAB7 (endosomes) (63). Ectopic expression in plants can provide insight into the subcellular localization of MpC1QBP as many eukaryotic localization signals are conserved (64). Moreover, plants are an attractive heterologous system for the study of plant virus-interacting proteins, as plant virus structural proteins are expressed and assembled in the plant host before encountering vector tissues.

In live plant epidermal cells, MpC1QBP-YFP signal was observed as punctate, motile bodies throughout the plant cytoplasm (Fig. 9A). Co-expression of MpC1QBP with the markers reveals this localization to also be multicompartmental (Fig. 9B-D), with patterns similar to what has been shown for C1QBP in human cells. Fluorescence from MpC1QBP-YFP co-localized with the mitochondrial matrix marker COX4-mCherry but was also observed independently in globular structures that came into contact with mitochondria (Fig. 9B). Further analysis indicated that MpC1QBP-YFP localized in a partially overlapping pattern with the Golgi-marker MAN49-(Fig. 9C) and the late-endosomal marker mCherry-RAB7 (Fig. 9D). Together, these data show that sequence signals for localization and possibly truncation of MpC1QBP are conserved between aphids and human C1QBP and that intracellular C1QBP docking sites are conserved in plants. We did not observe association of MpC1QBP-YFP with the plant plasma membrane, a pattern that was observed in our immunolocalization experiment in aphid guts (Fig. 8D-E).

### C1QBP functions as a negative regulator of PLRV transmission

To assess the function of C1QBP in PLRV transmission, we used a commercially available, small molecule inhibitor of HsC1QBP, M36. The inhibitor, identified in a pharmacophore screen by V. Yenugonda et al. (65), binds to HsC1QBP and inhibits its mitochondrial function in human glioma cells. *M. persicae* were exposed to a range of M36 concentrations (0, 50, 100 and 200 μM) supplied through sucrose diet sachets. After 48 hours of feeding on the inhibitor, aphids were moved to detached PLRV-infected or uninfected HNS leaves for a 24-hour acquisition access period (AAP) followed by a 72-hour inoculation access period (IAP) on 3-week-old potato (cv. Red Maria) seedlings. No differences in mortality were observed in the aphids as a result of the inhibitor treatments. The titer of PLRV in these recipient plants was then measured four weeks post inoculation (wpi) by double antibody sandwich enzyme-linked immunosorbent assay (DAS-ELISA). Results show that the percent of inoculated plants becoming systemically infected with virus was not statistically different when using aphids exposed to M36 compared to aphids that were not exposed to inhibitor (Fig. 10A, Table S3). However, plants that became systemically infected had higher titers of virus when aphids were exposed to concentrations of M36 ≥ 100 μM (Fig. 10B). When aphids were fed on diets containing 50 μM of M36, PLRV titer was variable and statistically similar to plants inoculated with aphids not exposed to M36. Conversely, plants inoculated with aphids fed on 200 μM of inhibitor exhibited a 1.63 fold-increase in virus titer that was statistically significant compared to the 0 and 50 μM M36 treatment conditions (*P* < 0.05). Exposure to 100 μM of inhibitor resulted in an intermediate phenotype, with a non-significant increase of titer relative to the control, but still less virus than in the 200 μM treatment (Fig. 10B). No obvious effects on mortality or visual effects on feeding behavior were observed at any of the concentrations of inhibitor.

**FIG 10.**
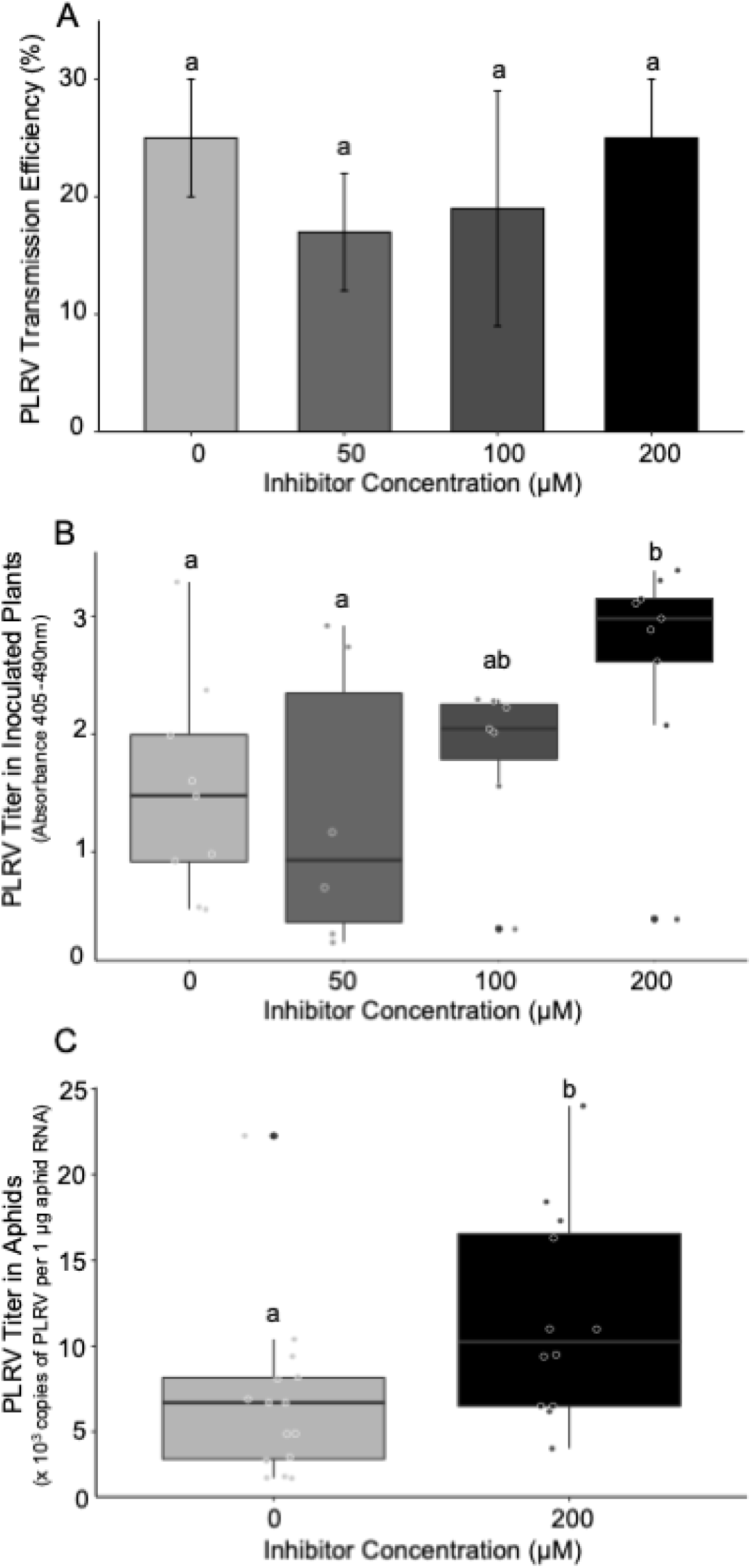
Transmission efficiency and virus titer in plants and aphids after aphid exposure to the M36 chemical inhibitor of C1QBP. *M. persicae* aphids were exposed to 0, 50, 100 and 200 μM of the M36 chemical inhibitor for 48 hours before transmitting PLRV from infected HNS leaves to potato seedlings. The number of infected potato plants and PLRV titer within those plants was assessed via DAS-ELISA after four weeks. (A) Bar graph of transmission efficiency of aphids exposed to the inhibitor, expressed as percent plants infected out of total plants inoculated (*n* = 51). Different letters indicate significantly different treatments (*P* < 0.05) by logistic regression analysis. Error bars represent ± one standard error. (B) Box plot of PLRV titer in infected potato plants (*n* = 5-9) inoculated by aphids exposed to various concentrations of inhibitor. Letters indicate significantly different treatments (*P* < 0.05) by a linear mixed effects ANOVA. (C) After 48 hours of aphid exposure to 0 or 200 μM of inhibitor and a 24-hour AAP on PLRV-infected HNS leaves, aphids were moved to sucrose diet for 3 days gut clearing and the copies of PLRV were quantified in each individual aphid (*n =* 12-15) by digital droplet PCR. Letters represent significantly different treatments by a one-tailed, unpaired Student’s *t-*test. B-C. Dots represent titer values colored by treatment. The thick black line indicates the median, with the box spanning the first and third quartiles. Lines reach out to the minimum and maximum values. Outliers are indicated with a black dot.

To test whether the observed increase in inoculated plant titer correlated with an increase in PLRV acquisition by the insects, aphids were exposed to 0 or 200 μM of inhibitor in sucrose diet for 48 hours and then moved to a PLRV-infected HNS leaf for 24-hour AAP (as in the transmission assay). After the AAP, the guts of aphids were cleared on fresh sucrose diet for 3 days to ensure detection of fully acquired and not ingested PLRV. The level of PLRV vRNA in individual aphids (*n*=12-15) was then measured by RT-digital drop PCR (ddPCR). A high percentage of aphids in both the control and inhibited treatments acquired virus (90% vs. 86%, data not shown), but on average, aphids exposed to 200 μM inhibitor acquired a significantly higher number of copies of PLRV (*P* = 0.015, 1.71-fold increase) compared to aphids fed on diet without M36. Together, these data suggest that C1QBP acts as a negative regulator of PLRV in the aphids, as inhibition of this protein by M36 leads to an increase in PLRV titer in the insects, and subsequently the aphid-inoculated plants.

## DISCUSSION

Optimization of several different extraction buffer conditions shows that no single extraction buffer tested captures all protein interactions formed between aphid proteins and PLRV. We encountered high variability between the three different trials of the experiment, even though there was significant capture of PLRV from viruliferous aphids in all biological replicates. This variability may be due to technical considerations, such as sub-optimal clarification of aphid homogenate increasing the capture of non-specific interactions, or biological reasons, such as the potentially transient nature of the interaction between some vector proteins and virus. Alternatively, variability in AP-MS experiments could be the result from true binding partners also having some high level of non-specific “stickiness” towards beads and/or antibody, thus making it hard to detect true signal above background (66). Other groups attempting to capture virus-vector protein complexes by AP have resorted to chemical cross-linking to preserve more transient interactions (19).

Transient interactions may be better captured by yeast two-hybrid where reconstitution of GAL4 stabilizes the interaction. This may explain why the vector proteins RR1 cuticle protein 5 and papilin isoform X1 were found to interact with luteovirids via Y2H and were enriched in our AP-MS experiments but considered lower confidence interactors. Interestingly, in the Y2H experiments, the RR1 cuticle protein from *S. graminum* directly interacted with the CP from luteovirid species transmitted by that aphid species but did not show interaction with PLRV CP, a virus *S. graminum* does not transmit, suggesting aphid-virus specificity or that the interaction is weaker/more transient between PLRV and *M. persicae*. This specificity phenotype is in-line with a previous study showing that vector competency of *S. graminum* was linked to specific protein isoforms that only differed by one amino acid from isoforms in non-competent lines of an F2 population (35). Taken together, these data suggest that even low confidence binding partners identified by AP-MS may be true direct binding partners of virus, and this technique can be complemented and the interactions confirmed by Y2H or other direct interaction methods. Moreover, this technique also identified at least one functionally relevant aphid protein interaction, MpC1QBP.

Addition of phosphatase inhibitor to the lysis buffer for the third AP-MS experiment allowed us to observe which interactions may be dependent on phosphorylation status. For instance, levels of cuticle protein 21 were significantly enriched by MS1 peak integration in APs from viruliferous aphids only when phosphatase inhibitor was added to the extraction buffer. However, RR1 cuticle protein-5, which directly interacted with the luteovirid CP in a virus-specific manner via Y2H, exhibited a trend of PLRV co-enrichment with or without phosphatase inhibitor, suggesting that this characteristic may only apply to certain classes of aphid cuticle proteins. Conversely, MpC1QBP and the viral non-structural polyprotein P1 were both significantly co-enriched with PLRV in all APs from viruliferous aphids except those where phosphatase inhibitor was added. These results suggest that interaction with PLRV may be contingent on phosphorylation of the vector proteins, protein complex partners, or the virion itself. Phosphorylation and acetylation of luteovirid capsid proteins have been previously shown (40, 67) although biological relevance of these post-translational modifications still remains to be determined.

Since PLRV is non-propagative in the aphid, the discovery of the P1 viral replication protein in PLRV protein complexes isolated from aphids was a surprising find. The aphids collected for AP were not subjected to gut-clearing prior to AP, thus our results could simply indicate capture of virion-P1 complexes from ingested sap in the aphid lumen. However, the ingestion of these complexes by the aphid raises the possibility that P1 may be internalized into aphids during virus acquisition. Indeed, P1 was found directly bound to purified TuYV virions using chemical cross-linking mass spectrometry (68). Furthermore, the PLRV P1 protein suppresses aphid-induced jasmonic acid signaling in *N. benthamiana* indicating a role for P1 in vector manipulation (69). One hypothesis our data generates is that P1 protein-PLRV virion complexes that are acquired from infected plants. These P1-virion complexes may circulate through the aphid and be inoculated into new host plants together, which may help nascent PLRV infections to become established in new host plants. Immunolocalization of the P1 polyprotein within aphid cells would support this hypothesis.

While cuticle proteins have recently been identified as putative receptors for nonpersistent viruses that bind cuticle-lined aphid mouthparts (13, 15), the exact role of cuticle proteins in the transmission of circulative viruses remains elusive. Cuticular proteins are frequently identified in vector-plant virus interaction assays (23, 26). For example, out of six western flower thrips proteins found to interact with TSWV in a study by I. E, Badillo-Vargas et al. (23), half were annotated as cuticle proteins. Genes coding for cuticle proteins were also the largest group found to be responsive to the tospovirus *Tomato spotted wilt virus* (TSWV), with the majority of these genes being downregulated in the larval and prepupal stages of the insect when TSWV is acquired (30). An aphid cuticle protein was also identified as binding *Beet western yellows virus* (BWYV, Genus: *Polerovirus*) *in vitro*, another luteovirid that is transmitted by *M. persicae* (26). Authors hypothesized that the interaction between luteovirids and cuticle proteins could represent an ancient, less efficient mode of virus transmission in aphids. In addition to comprising the exoskeleton of the insect, cuticle proteins are present in the foregut and hindgut in many hemipterans, locations within the alimentary canal where luteovirids are either acquired or possibly excluded from moving into gut cells. Determining where viral interaction with cuticular proteins occurs can be the key to understanding their function. For instance, the cuticle protein CPR1 from the insect vector *Laodelphax striatellus* was found to interact with the tenuivirus *Rice stripe virus* and co-localize with virion in insect hemocytes (14). Knockdown of CPR1 led to decreased virus accumulation in the hemolymph and salivary glands, while transmission to plants was reduced. The authors concluded that CPR1 might bind and stabilize RSV in the hemolymph to protect it from degradation or facilitate passage to the salivary glands. Further tissue-specific interaction or localization studies with the PLRV-interacting cuticle proteins identified in our study as well as functional assays could shed light on the role these proteins play in the transmission or PLRV, or poleroviruses in general.

Complement component 1 Q subcomponent-binding protein (C1QBP) was found to be the most enriched vector protein in α-PLRV APs from viruliferous aphids compared to non-viruliferous controls in the absence of phosphatase inhibitor. A protein orthologue in *S. graminum* directly interacted in Y2H assays with the CP domain of multiple virus species from the family *Luteoviridae*, including PLRV, indicating a more conserved role for C1QBP in circulative transmission. The C1QBP protein is evolutionary conserved; with sequences being found in other distantly related eukaryotic species (70). Phylogenetic analysis indicates that C1QBP from aphids is more closely related to human C1QBP that has evolved multiple functions and is less related to C1QBP in plants and yeast, which so far has only been shown to function in oxidative phosphorylation (56).

The putative function of human C1QBP has been extensively studied due to the finding that its expression is highly elevated in cancer cells (54), and bialleic mutations of *C1QBP* in the human population are associated with a mitochondrial disorder that leads to cardiomyopathy (71). Structurally, C1QBP forms a donut-shaped homotrimer (72). Its conserved eukaryotic function is as a molecular chaperone involved in ribosomal biogenesis in mitochondria (56, 71). However, in humans, the protein is multifunctional, having roles in innate and adaptive immunity, inflammation, infection, apoptosis, pre-mRNA splicing, macrophage cell adhesion and cancer cell metastasis (73–78). C1QBP was first identified as a binding partner of C1q (79), a component of the complement pathway in human blood which binds antibody-antigen complexes and activates the complement system (80). However, HsC1QBP also binds directly to capsid proteins of numerous viruses, including rubella (81), human immunodeficiency virus type 1 (82), and herpes simplex virus 1 (83). Collectively, our interaction experiments indicate the ability of C1QBP to bind viruses may also be evolutionarily conserved in insects.

During circulative transmission, luteovirid virions are acquired into aphid gut cells via receptor-mediated endocytosis and trafficked across the cell in clathrin-coated vesicles that can be tubular in structure (5). A. Garret et al. (6) shows that virions are sometimes observed in lysosomes or multivesicular bodies (MVBs), but the exact role these organelles have in transmission still remains uncharacterized. It is also unknown whether virus-containing vesicles contact the mitochondria, the primary location of C1QBP in human cells. Therefore, based on what has been published about luteovirid trafficking, it is difficult to say in which subcellular compartment MpC1QBP and PLRV are interacting in the aphid gut epithelial cell cytoplasm, and whether this localization is necessary for the normal transcytosis of virions. Considering that live-cell imaging studies of eukaryotic cells have shown that most organelles are not static, discrete compartments but rather dynamic entities that actively associate with one another (84), the hypothesis that virus-containing vesicles do come in contact with other organelles as they traverse the aphid cell, is plausible. The development of robust subcellular markers in aphids is definitely needed to determine conclusively what subcellular compartments MpC1QBP and PLRV occupy.

As for the function of MpC1QBP in luteovirid transmission, we can turn to the known roles of C1QBP in human innate immunity for clues. Virus binding to vector proteins may be an effort by the virus to hijack vector machinery to facilitate its transmission or an effort by the insect to defend against the virus. On the surface of cells, HsC1QBP can act as a receptor for viral entry into cells (85–87). Infection of human cells with Sendai virus leads to a re-localization of C1QBP to the mitochondria, where it dampens the RIG-I and MDA5-dependent (MAVS) antiviral defense response by binding key signal transduction molecules, which results in the promotion of viral replication (88). In the case of HsC1QBP binding to rubella virus, it was found that inhibition of C1QBP slows microtubule-directed redistribution of mitochondria and reduces rubella virus replication (89). This led the authors to hypothesize that rubella capsid protein binding to C1QBP helps traffic mitochondria near sites of rubella virus replication to help supply energy. Thus, in human cells, some viruses have evolved to commandeer the function of C1QBP to support propagation. In the case of MpC1QBP interaction with PLRV, exposure to the C1QBP chemical inhibitor M36 had the opposite effect and lead to an increase of PLRV titer in aphids. For a non-propagative virus, ways this may be achieved include an increase in the rate of virus acquisition or an enhanced stability of the acquired virus in aphid tissues. Our data supports the role of MpC1QBP as a negative regulator of PLRV accumulation in the aphid. Restricting the amount of virus in the insect would be consistent with the role of a defense protein. Therefore, it is possible that the innate immune function of C1QBP is conserved in aphids.

However, because PLRV is non-propagative in its aphid vector, we would not expect the insects to launch an immune response to the virus. Nevertheless, this does not preclude aphids from recognizing luteovirids as a pathogen based on their icosahedral geometry typical of viruses. Therefore, we hypothesize that MpC1QBP recognizes the pathogen-associated molecular pattern (PAMP) of the luteovirid capsid and activates or participates in some immune response that leads to the reduced transmission of PLRV. If MpC1QBP does act as a sentinel for viral invaders, then this is in line with our shotgun proteomics data that shows MpC1QBP is expressed in non-viruliferous aphids, and its level is unchanged in response to PLRV. It is unknown what innate immune pathway MpC1QBP would be a part of, as insects do not have the classical complement pathway of higher organisms, although proteins with C1q-like domains have been detected in insects, including the pea aphid (90). Aphids also lack the immunodeficiency (IMD) pathway common in other insects, though homologues for the constituents of other conserved innate immune pathways, such as JAK-STAT and Toll, have been found in aphid genomes (91). It should be noted that, even though our imaging was limited to the gut, our interaction and functional studies were conducted using whole aphids. Therefore, the inhibitory function of MpC1QBP in other parts of the aphid involved in circulative transmission, such as the hemolymph (insect blood) and the accessory salivary glands, cannot be distinguished.

Comparing the acquisition data to the transmission data for inhibition of MpC1QBP, it is interesting that an increase in the amount of virus acquired by the insects did not necessarily correlate with an increase in percent transmission. In our experiments, the consequence of inhibited aphids acquiring more virus does not seem to be more transmission, but rather higher virus titer in the plants that do become infected. A possible explanation is that, in the conditions used for our experiment, a high percentage of aphids in the control treatment still acquired enough virus to launch a systemic infection in the plant, but aphids in the inhibitor treatment may have injected more initial virus inoculum, resulting in higher systemic titer in the infected plants. This observation may explain why PLRV has yet to evolve some mechanism to avoid binding C1QBP since, even though it is a negative regulator, C1QBP does not adversely affect transmission rate and ultimately, virus spread. In fact, as obligate biotrophs, viruses must carefully regulate their replication to not kill their host. Therefore, the role of C1QBP in reducing virus titer in the insect and plant may not only protect the insect from too much virus, but also indirectly benefit or protect the plant host.

A limitation of our study is our inability to measure how much M36 entered into aphid cells. Importantly, no aphid mortality was observed during feeding on M36 inhibitor. Functional genomics in aphids is a major challenge in the field. Gene silencing is incomplete in aphids, and the timing of silencing experiments are complex when coupled to studying phenotypes in virus transmission. Thus, working with inhibitors that have well characterized modes of action in other animals is an attractive alternative that allows us to probe protein function in virus transmission. However, inhibition of C1QBP by the M36 inhibitor could have had other unknown effects on the aphid. For example, if M36 increased aphid feeding, we would have also observed an increase in virus acquisition and transmission. Additional experiments, such as electrical penetration graph (EPG) assays, would be necessary to discern between these two possible scenarios.

## MATERIALS AND METHODS

### Plants, Insects and Virus

A parthenogenetically-reproducing clone of *M. persicae* Sulzer (red strain) (92) was maintained on *Physalis floridana* at a temperature of 20°C with a photoperiod of 16-hour day/8-hour night. Colonies of *Schizaphis graminum* genotypes SC and F (93) were reared on oats (cv. Coast Black) at 20°C in 24-hour light conditions. A cDNA infectious clone of *Potato leafroll virus* (PLRV) developed by L. F. Franco-Lara et al. (94) was used to inoculate hairy nightshade (*Solanum saccharoides*, HNS) via agroinfiltration (95), which was used as a source of virus inoculum for all subsequent experiments. PLRV-infected HNS plants were used ∼3-4 weeks post-infiltration when the disease symptom of interveinal chlorosis was visible on most leaves. Hairy nightshade plants were grown in growth chambers with a set temperature of 20°C and a photoperiod of 16-hour day/8-hour night. *Nicotiana benthamiana* used for microscopy and western analysis were grown under the same conditions described in (96).

### Extraction of virus-vector protein complexes

*M. persicae* were allowed a 48-hour acquisition access period (AAP) on PLRV-infected or healthy HNS under 24-hour light conditions. After collection, aphids were stored at - 80°C for further use. Aphids pooled from 1 to 3 separate collections were cryogenically ground using a Mixer Mill MM 400 (Retsch). Approximately 2-4 g of tissue were ground in 10 mL MM 400 metal canisters with one 12 mm steel ball for 6 sets of 3 minutes at a vibrational frequency of 30 Hz with a five-minute liquid nitrogen cooling period in between grinding cycles. After cryogenic lysis, the resulting powder was stored at −80°C for far-western analysis and affinity purification.

For optimization of lysis buffer conditions by far-western blotting, cryogenically lysed powder from aphids fed on healthy HNS was split equally and solubilized in the following buffers on ice: filter sterilized CHAPS buffer = 1X phosphate buffered saline (PBS, pH7.4), 40 mM 3-[(3-cholamidopropyl)dimethylammonio]-1-propanesulfonate (CHAPS), 10 mM calcium chloride; HEPES buffer = 50 mM HEPES-KOH (pH 7.4), 110 mM KOAc, 2 mM MgCl_2_, 0.4% TritonX-100; TBT buffer = 50 mM HEPES-KOH (pH 7.4), 200 mM tris(hydroxymethyl)aminomethane (Tris, pH7.5), 110 mM KOAc, 2 mM MgCl_2_, 0.4% TritonX-100, 0.1% Tween-20, 350 mM sodium chloride; TRIS buffer = 50 mM Tris (pH7.5), 150 mM sodium chloride, 0.4% TritonX-100; SDS = 50 mM Tris (pH 6.8), 2.5 % Sodium dodecyl sulfate, 10% glycerol. All buffers were supplemented with 0.5 mM phenylmethylsulfonyl fluoride (PMSF) (Sigma-Aldrich) and a 1:100 (v/v) dilution of Halt™ EDTA-free protease inhibitor cocktail (PI) (Pierce). One milliliter of buffer was added to 200 milligrams of powder and incubated on ice for 10 minutes, with vigorous vortexing every 2 minutes. Samples were then centrifuged at 16,100 x *g* for 10 minutes at 4 °C. Supernatant was moved to a fresh tube, photographed and stored on ice for far-western analysis.

### Far-western analysis

The supernatants from the different lysis buffer extractions described above were diluted 1:4 in 4x Laemmli sample buffer (Bio-Rad) supplemented with beta-mercaptoethanol following manufacture’s instructions. Samples were incubated at 95°C for 10 minutes followed by centrifugation at 16,100 x *g* for 1 minute at room temperature. Forty μL of the CHAPS, HEPES and TBT extractions and 20 μL from the TRIS and SDS samples were separated side-by-side on two separate pre-cast 4-20% TGX gradient gels (Bio-Rad) by SDS-PAGE and proteins transferred to nitrocellulose following the protocol described in (40). Blots were then blocked for 15 minutes in TBS with 2% (v/v) Tween 20 (-T) followed by an overnight block in 5% non-fat dried milk in TBS-T (0.1%), gently rocking at room temperature. Blots were washed briefly in 1x TBS-T (0.1%) and incubated with 10 mL of 1x TBS-T (0.1%) supplemented with 8.6 μg of gradient purified PLRV (97) or TBS-T (0.1%) without virus (negative control) overnight at 4°C, with gentle rotation. The next day, blots were washed four times in TBS-T (0.1%) for 10 minutes each at room temperature and then incubated for 1 hour at room temperature with a 1:2500 dilution (TBS-T 0.1% supplemented with 0.1% BSA) of the PLRV primary antibody described in (42). Antibody was removed and blots washed again as described above. A 1:2500 dilution of alkaline phosphatase conjugated α-rabbit secondary antibody (Sigma) in TBS-T 0.1% supplemented with 0.1% BSA was applied and blots incubated for 2 hours at room temperature followed by the washing procedure described above with the addition of a final 10 min, room temperature wash in TBS. Blots were incubated with 1-Step™ NBT/BCIP substrate solution (Thermo Scientific) for exactly 1 minute 47 seconds and washed quickly with deionized water to stop development.

### Affinity purification

For each biological replicate, four grams of cryogenically-lysed *M. persicae* tissue (viruliferous or non-viruliferous) was solubilized in 20 mL of the TRIS extraction buffer described above. For APs with the phosphatase inhibitor, the protease inhibitor cocktail was replaced with the same amount of Halt™ Protease and phosphatase inhibitor single-use cocktail, EDTA-free (Pierce). Total protein was extracted following the lysis protocol outlined in (40) with a few exceptions. Samples were incubated at 4°C for ∼20 minutes on ice with occasional vortexing. Tissue was then homogenized in buffer, on ice, with a Polytron (Brinkmann Instruments) for two, 10-second pulses at setting “2” separated by a 30-second incubation on ice. Lysates were then rotated at 4°C for 10 minutes. The resulting homogenate was moved to a glass Corex centrifuge tube and centrifuged for 10 min at 4°C in a Beckman Avanti J-25 I centrifuge in a JA-20 rotor to remove cell debris. The speed of centrifugation varied for each AP experiment. The April samples were centrifuged twice at 3500 rpm (1,480 x *g*) and once at 6000 rpm (4,355 x *g*). For the January dataset, biological replicates 1 (JMPH1 and JMPW1) were centrifuged once at 6000 rpm (4,355 x *g*). All other January biological replicates and APs from the Phosphatase Inhibitor dataset were centrifuged once at 8000 rpm (7,740 x *g*). Each independent dataset consisted of three biological replicate APs for viruliferous and non-viruliferous controls. Biological replicates were separate collections of pooled aphids fed on a group of systemically infected or healthy HNS plants.

For all APs analyzed by mass spectrometry, after centrifugation, the aqueous layer was removed and diluted 1:2 in fresh TRIS extraction buffer on ice. Five milligrams of α-PLRV conjugated M-270 epoxy Dynabeads™ (Invitrogen) were rotated with 10 mL of the diluted aphid homogenate for 1 hour at 4°C and beads washed with TRIS extraction buffer as described in (40). Final wash buffer was completely removed from magnetic beads and captured proteins were subjected to on-bead reduction, alkylation and trypsin digestion following the protocol outlined in (42). Smaller volume APs (1 mg beads per 2 mL of diluted homogenate) were done in parallel for each of the three independent AP datasets and analyzed by Western analysis to gauge the enrichment of PLRV structural proteins (40).

### Mass spectrometry and data analysis

For AP samples, tryptic peptides were reconstituted and analyzed on an Orbitrap Fusion Tribrid mass spectrometer (Thermo Scientific) following the parameters described in (42). Each affinity purification sample was analyzed a total of three times. For two of the analytical replicates, the fragment ions were analyzed in the linear trap using “rapid” scan rate. For the third analytical replicate, fragment ions were analyzed in the Orbitrap. Two analytical replicates for the April dataset were analyzed months prior to the January and Phosphatase Inhibitor datasets but a third analytical replicate from the April dataset (MS^2^ captured in the linear trap) was analyzed alongside all samples from the two other datasets.

Protein identification of resulting mass spectrometry data from APs was conducted as described in (42) with a few exceptions. The protein search database was generated from amino acid sequences corresponding to all coding gene sequences from the *Myzus persicae* G006 genome assembly v1.0 obtained from AphidBase (https://bipaa.genouest.org/sp/myzus_persicae_g006/) on the Bioinformatics Platform for Agroecosystems Arthropods (BIPAA) website prior to its formal release as well as sequences from all known species of *Luteoviridae* and common mammalian AP contaminant proteins downloaded from NCBI. Mascot search parameters were changed slightly from those outlined in (42) so that the mass measurement accuracy was set at 0.5 Da or 0.02 Da for fragment ions for analytical replicates where MS^2^ spectra were collected in the linear trap or Orbitrap, respectively.

Mascot search results for APs were imported into Scaffold-Q+ version 4.8.9 for label free quantification by spectral counting using the protein cluster feature with the following identification filter thresholds: two-peptide minimum and a 1% false discovery rate on both the peptide and protein level. Vector proteins with significant overlapping shared-peptide evidence that were identified as a cluster were treated as a proxy for a single identification, and total spectral counts were calculated on the level of the whole cluster with the protein reference number listed corresponding to the protein with the most unique peptides assigned in the cluster. Precursor ion (MS1) peak areas for peptides corresponding to selected proteins (Table S2) were measured using Skyline (98). Statistical analysis of AP-MS datasets was done using the T-test and ANOVA features in Scaffold-Q+ and Significance Analysis of Interactome (SAINT) Express probability scoring through the crapome.org web interface (47).

Shotgun MS analysis comparing C1QBP protein levels between viruliferous and non-viruliferous aphids reared on physalis was conducted and data analyzed exactly as described in (51) for comparing proteins differentially regulated in physalis- and turnip-reared *M. persicae*.

### Yeast-two-hybrid

Total mRNA was purified from a pool of *S. graminum* aphids, genotype F2-A3, an efficient vector of CYDV-RPV (93), reared on RPV-infected plants using the RNeasy Plant extraction kit (Qiagen) followed by capture with oligo-dT magnetic-beads (PolyATtract(R) mRNA Isolation Systems, Promega). A full-body cDNA library was constructed from the *S. gramiunum* RNA in the BD Matchmaker AD cloning vector (pGADT7-Rec, BD Biosciences Clontech) following the manufacturer’s instructions, and transformed in yeast strain AH109. The transformed cells were plated on synthetic dropout (SD) selection media lacking leucine (SD-Leu). Transformants were recovered and pooled in freezing medium as 1 mL aliquots with a concentration above 2 x 10^7^ cells/mL were stored at −80°C. The coat protein (CP) and readthrough domain (RTD) genes of BYDV-PAV, CYDV-RPV and PLRV were cloned separately into a DNA-BD fusion vector (pGBKT7, BD Biosciences Clontech) and transformed in yeast strain Y187.

The *S. gramiunum* cDNA library was screened by separately mating with the Y187 luteovirid coat protein strains following manufacture guidelines. Briefly, one aliquot of AH109-library (≥ 2 x 10 cells) was combined with each Y187-bait culture and incubated for 24 hours at 30°C in 45 mL 2x YPDA supplemented with Kanamycin (50 μg/mL, Kan). The mating mixture was centrifuged and resuspended in 10 mL of 0.5x YPDA/Kan (50 μg/mL) and plated on SD selection media lacking tryptophan, leucine, and histidine (SD-Trp/-Leu/-His). Positive clones (yeast colonies growing on SD/-Trp/-Leu/-His media) were subcultured on SD/-Trp/-Leu/-His/-Ade containing X-α-Gal (0.4 mg/mL) to detect strong, positive interactions. Plasmid was extracted from positive clones, re-transformed into DH5α *E. coli* cells by heat shock, and cloned cDNA sequences identified by Sanger sequencing using pGAD Rec T7-specific primers. The resulting sequences were compared against EMBL and GenBank sequences using BLAST. The ability of candidate proteins from the *S. graminum* cDNA library to interact with PAV, RPV or PLRV CP or RTD proteins was confirmed using the co-transformation approach. Co-transformation with DNA-BD fusions of human lamin C and the murine p53 was used as negative controls.

### Sequence alignment and phylogenetic analysis

Orthologous protein sequences for MpC1QBP were identified using NCBI BLAST, which calculates percent identity and similarity. Sequence alignments were generated using the Clustal Omega Multiple Sequence Alignment web interface (https://www.ebi.ac.uk/Tools/msa/clustalo/) and visualized with BoxShade (https://embnet.vital-it.ch/software/BOX_form.html). Phylogenetic analysis was performed using the one-click mode of the Phylogeny.fr web tool (https://www.phylogeny.fr/) without Gblocks curation. NCBI or citrusgreening.org accession numbers for protein sequences used are: XP_026823488.1 (*Rhopalosiphum maidis*), XP_025194941.1 (*Melanaphis sacchari*), XP_027842714.1 (*Aphis gossypii*), NP_001233078.1 (*Acyrthosiphon pisum*), XP_015380335.1 (*Diuraphis noxia*), XP_025410594.1 (*Sipha flava*), DcitrP057835.1.1 (*Diaphorina citri*), RZF37806.1 (*Laodelphax striatellus*), XP_014240636.1 (*Cimex lectularius*), XP_018897713.1 (*Bemisia tabaci*), XP_012542024.1 (*Monomorium pharaonis*), ETN58839.1 (*Anopheles darling*), NP_611243.1 (*Drosophila melanogaster*), NP_001203.1 (*Homo sapiens*), NP_001320214.1 (*Arabidopsis thaliana*), KZV10545.1 (*Saccharomyces cerevisiae*).

### Immunolocalization in aphids

Adult and late instar *M. persicae* were caged on detached PLRV-infected or uninfected HNS leaves for 48 hours in 24-hour light conditions. Guts from aphids exposed and not exposed to PLRV were dissected into PBS. Guts were fixed in 4% formaldehyde for 30-45 minutes before being permeabilized with 0.1% (v/v) Triton X-100 for 2 hours. Guts were washed with PBS-T (PBS, 0.5% (v/v) Tween 20) 3 times and blocked for 2-3 hours in PBS-T, 1% (m/v) BSA. After blocking, guts were incubated with cross-absorbed α-PLRV diluted 1:1000 in blocking buffer (PBS-T, 1% BSA), 1:50 polyclonal antibody against full-length human C1QBP (α-HsC1QBP) derived from mouse (Sigma-Aldrich), or both overnight at 4°C. Guts were washed 5 times in PBS-T and incubated in secondary antibody, 1:500 donkey α-rabbit-Cy3 (Millipore Sigma), or 1:250 donkey α-mouse-Cy2 (Jackson ImmunoResearch), diluted in blocking buffer for 1 hour. Guts were washed 5 times in PBS-T and then mounted on slides in Flouromount plus 4’,6-diamidino-2-phenylindole nuclear stain (DAPI, Southern Biotech) and sealed with a cover slip. Guts were visualized with a Leica TCS SP5 laser scanning confocal microscope. Cy2 was excited with the 488-nm line of a multiline Argon laser with emission spectra collected by a photomultiplier tuber (PMT) detector in the range of 545-550 nm (Fig. 8B, D) or a hybrid detector (HyD) in the range of 500-530 nm (Fig. 8C, E). Cy3 was excited with the 561-nm line of multiline Argon laser with emission spectra collected by a HyD in the range of 604-631 nm. DAPI was excited by a 405-nm ultraviolet laser with emission spectra collected by a HyD in the range of 445-479nm. All scans were conducted sequentially with line averaging of 6 or 8. Non-viruliferous guts stained with α-C1QBP and α-PLRV as well as viruliferous guts stained with only secondary antibodies were used as negative controls. The experiment was repeated independently twice.

### Ectopic expression in plants

The full-length coding sequence of *C1QBP* without a stop codon was amplified from *M. persicae* cDNA with Phusion™ High-Fidelity DNA polymerase (Thermo Scientific) following manufacture’s guidelines with the following primers, Forward: 5’-GGGGACAAGTTTGTACAAAAAAGCAGGCTTA<u>ATGAATACTTTAATCAGATCG</u> and Reverse: 5’ – GGGGACCACTTTGTACAAGAAAGCTGGGTG<u>AAAAAATTTCCTTAAATCA</u> – 3’. Underlined nucleotides correspond to the MpC1QBP sequence fused to attB Gateway™ cloning sites. The resulting amplicon was cloned into the pEarleyGate 101 destination vector (99) using Gateway™ technology (Invitrogen) as described in (96) to create the p35s:MpC1QBP-YFP construct. This construct was transiently expressed via agroinfiltration with the organelle markers, COX4-mCherry, Man49-mCherry or mCherry-Rab7 (63) in *N. benthamiana* epidermal cells and imaged 3 days post-infiltration with a Leica TCS SP5 laser scanning confocal microscope as described in (96).

### Chemical Inhibition of C1QBP

*M. persicae* aphids were collected and starved for 1-2 hours before being placed on artificial sucrose diet (51) containing 0, 50, 100, or 200 μM of a small molecular inhibitor of human C1QBP (65) resuspended in water, diluted in sucrose diet and 0.1% dimethyl sulfoxide (DMSO) in a membrane feeding sachet. After 48 hours of feeding on the inhibitor, aphids were moved to detached PLRV-infected or uninfected HNS leaves for a 24-hour AAP. Then, aphids were moved to 3-week-old potato cv. Red Maria seedlings, 5 aphids per plant, 12-15 plants per treatment, for a 72-hour inoculation access period (IAP). After the IAP, aphids were removed with an application of pymetrozine (Endeavor) and bifenthrin (Talstar P). Two- and four-weeks post inoculation potato plants were assessed for systemic PLRV infection and titer. Four leaf discs were taken from the youngest fully emerged leaf on the apical stem of the potato plants and used for double antibody sandwich enzyme-linked immunosorbent assays (DAS-ELISA) with a commercially available PLRV antibody kit (Agdia). The transmission assay was repeated independently three times.

For virus acquisition experiments, aphids were fed on sucrose diets containing 0 or 200 μM of inhibitor for 48 hours and then moved to a PLRV-infected HNS leaf for 24-hour AAP (as in the transmission assay). After the AAP, aphids were moved to fresh sucrose diet for 3 days gut clearing to remove any residual PLRV in their gut lumen so any detected PLRV was acquired across the midgut of the insects. After gut-clearing, individual aphids (*n* = 12-15) were collected and flash frozen at −80 °C. RNA was extracted from the aphids using the RNeasy Mini Kit (Qiagen) and cDNA was synthesized from 0.1 ug of aphid RNA using the iScript cDNA Synthesis Kit (Bio-Rad). PLRV titer in undiluted cDNA was quantified by digital drop PCR using EvaGreen Supermix and the QX100 droplet digital PCR system as previously described (51).

### Statistical Analysis

PLRV transmission efficiency after aphid exposure to the inhibitor was analyzed using logistic regression. The model predicts whether an inoculated plant will become infected based on different predictors. A one-tailed likelihood ratio test showed that experiment could be removed from the model (*P* = 0.578). Therefore, the model has the sole predictor: treatment. The full model, model diagnostics, test statistics, and *P*-values are reported in Table S3. PLRV titer in inoculated plants was analyzed with a linear mixed effects analysis of variance using Satterthwaite’s method. Inhibitor concentration was used as a fixed effect and experiment as a random effect. Linear contrasts were performed using Dunnett’s test. For PLRV titer in the insects, the differences in titer between the two treatments was compared using a one-side, unpaired Student’s *t* test. Letters represent statistically different groups (*P* < 0.05). All error bars represent ± one standard error. All indicated analyses were performed in R version 3.6.3 (https://www.r-project.org/).

### MS Data Availability

Heck, M. (2020) Isolation of aphid-*Potato leafroll virus* (PLRV) proteins complexes from viruliferous insects using affinity purification-mass spectrometry. Available from ProteomeXchange.org using the project identifier PXD022167.

## ACKNOWLEDGMENTS

The authors would like to thank the following: former technicians Dawn Smith, Ana Rita Rebelo, and Rogerio Santos for their help in the care and collection of insects; technicians Maria G. Gutierrez, Myah Frostclapp, and the BTI and Cornell greenhouse staff for plant care; Jason Ingram (USDA-ARS) for application of pesticides. John Flaherty and the IT staff of Cornell University’s Biotechnology Department for help with Mascot support; Maria Harrison (BTI) and Sergey Ivanov for use of their mCherry organelle marker constructs; Mamta Srivastava (BTI, Plant Cell Imaging Center) for technical help with imaging and microscopy software.

This work was supported by USDA ARS CRIS project 8062-22410-006-00-D, USDA NIFA project 8062-22410-006-15-A. J.R.W. is supported by an NSF Graduate Research Fellowship (DGE-1650441) and a NIFA Predoctoral Fellowship (2019-67011-29610).

## SUPPLEMENTARY DATA FOOTNOTES

**Table S1: Viral and vector proteins found to be ≥2-fold enriched in PLRV affinity purifications from viruliferous aphids in one or ore independent AP-MS experiments**

*^a^*Prey proteins (or clusters) are grouped based on the number and significance of AP-MS dataset(s) they were found to be enriched ≥2-fold or present/absent (+/-) in APs from viruliferous aphids compared to respective negative controls.

*^b^*Column indicates the dataset(s) prey proteins (or clusters) were found to be significantly enriched in APs from viruliferous aphids compared to their respective non-viruliferous negative controls.

*^c^*Protein accession corresponding to the viral and *Myzus persicae* protein sequences within our MS search database that was based on version 1.0 of the *M. persicae* (clone G006) genome assembly obtained from BIPAA AphidBase and viral sequences from NCBI. Bracketed integers denote the identification of a protein cluster and the number of proteins belonging to that cluster. Integers in parentheses indicate the number of additional proteins that share the same exact peptides identified by MS.

*^d^*Functional annotation of prey protein sequences obtained from NCBI p rotein BLAST.

*^e^*Protein identifier for corresponding reference sequence in NCBI.

*^f^*Protein molecular weight based on amino acid sequence.

*^g^*TRUE denotes a protein cluster or protein that shares some but not all peptide sequences with other proteins identified.

*^h^*Highlights the AP category the indicated prey protein (or cluster) was determined to be significantly enriched (p-value < 0.05) by Student T-test in Scaffold Q+ using total spectral counts.

*^i^*Highlights the AP category the indicated prey protein (or cluster) was determined to be significantly enriched (p-value < 0.05) by ANOVA in Scaffold Q+ using total spectral counts.

*^k^*Quantitative MS data representing the set of APs performed and analyzed in April 2015. Biological replicates (n=3) corresponding to PLRV APs from non-viruliferous (N) and viruliferous (V) *M. persicae* are denoted as AMPH and AMPW, respectively, in this table.

*^m^*Quantitative MS data representing the set of APs performed and analyzed in January 2016 where phosphatase inhibitor cocktail was added to the AP lysis buffer. Biological replicates (n=3) corresponding to PLRV APs from non-viruliferous (N) and viruliferous (V) *M. persicae* are denoted as PhoIn_H and PhoIn_W, respectively, in this table.

*^p-q^*Average value of total spectral counts for PLRV APs from non-viruliferous and viruliferous *M. persicae* within the indicated dataset.

*^r^*Fold enrichment calculation based on the ratio of the average total spectral counts detected in APs from viruliferous aphids to non-viruliferous negative controls within the indicated dataset. (+/− denotes presences/absence; nd = not detected).

*^s^*T-test p-value calculated in Scaffold Q+ comparing the average total spectral counts detected in APs from viruliferous aphids to non-viruliferous negative controls within a dataset.

*^t^*Significance Analysis of INTeractome (SAINT) probability score indicating interaction confidence of each prey protein in APs from viruliferous aphids compared to non-viruliferous negative controls within a dataset. High confidence interaction scores are ³0.8, medium confidence interaction score are between 0.5 and 0.79, low or no confidence scores are ≤ 0.5. Scores in red indicate the SAINT probability of interaction when the total spectral counts detected in each of the indicated APs from viruliferous aphids were compared to all non-viruliferous negative controls, including ones from the other two datasets.

**Table S2. Peptides measured by MS1 peak integration**

*^a^*NCBI reference number corresponding to protein sequences with peptide spectral matches in our AP samples.

*^b^*Protein symbol of viral and vector proteins whose levels were analyzed by MS1 peak integration, including immunoglobulin G (IgG) that was coupled to the magnetic beads.

*^c^*Amino acid sequence of peptide ions deduced from MS2 fragmentation that were used to quantify protein levels in AP samples by MS1 quantification. The residue position of the start and end of each peptide within the corresponding sequence in our protein database is shown in brackets. The amino acid residues before and after trypsin cleavage are given as a reference. Missed 1 indicates a missed cleavage site. Modified residues are underlined and bold faced.

*^e^*Average peptide retention time given in minutes.

*^f^*Charge state of peptide precursor ion analyzed.

*^g^*Sum of the integrated MS1 peak area for all precursor isotope ions ([M]+[M+1]+[M+2]) minus background measured in each analytical replicate of an affinity purification (AP) sample using Skyline MS1 full-scan filtering and manual peak boundary refinement. The data is segregated into the three independent AP-MS experiments: the April, January and Phophatase Inhibitor datasets.

*^h^*Sample names for non-viruliferous (NV) and viruliferous (V) AP analytical replicates written as AP_biological replicate number_analytical replicate number. The corresponding raw MS file available via ProteomeXchange with identifier PXD022167 is given in brackets.

**Table S3. Logistic Regression Analysis of PLRV Transmission by M. Persicae Aphids after Exposure to the C1QBP Chemical Inhibitor**

*^a^*Model output is the categorical variable “InfectionState” with levels 0 = uninfected and 1 = PLRV-infected indicating whether the inoculated plant became systemically infected.

*^b^*“Treatment” is a quantitative variable indicating the concentration of the chemical inhibitor delivered to the aphids.

*^c^*“Exp” is a categorical variable representing the different trials of the experiment. Abbreviations: *SE*, standard error; *df,* degrees of freedom; Ho, null hypothesis

## REFERENCES

1. Nault LR. 1997. Arthropod transmission of plant viruses: A new synthesis. Annals of the Entomological Society of America 90:521–541.

2. Gray SM, Banerjee N. 1999. Mechanisms of arthropod transmission of plant and animal viruses. Microbiol Mol Biol Rev 63:128–148.

3. Brault V, Herrbach E, Reinbold C. 2007. Electron microscopy studies on luteovirid transmission by aphids. Micron 38:302–312.

4. Gildow FE. 1987. Virus-membrane interactions involved in circulative transmission of luteoviruses by aphids, p 93–120. *In* Harris KF (ed), Current Topics in Vector Research, vol 4. Springer-Verlag, New York.

5. Gildow FE. 1993. Evidence for receptor-mediated endocytosis regulating luteovirus acquisition by aphids. Phytopathology 83:270–277.

6. Garret A, Kerlan C, Thomas D. 1996. Ultrastructural study of acquisition and retention of potato leafroll luteovirus in the alimentary canal of its aphid vector, *Myzus persicae* Sulz. Arch Virol 141:1279–1292.

7. Garret A, Kerlan C, Thomas D. 1993. The intestine is a site of passage for potato leafroll virus from the gut lumen into the haemocoel in the aphid vector, *Myzus persicae* Sulz. Arch Virol 131:377–392.

8. Peiffer ML, Gildow FE, Gray SM. 1997. Two distinct mechanisms regulate luteovirus transmission efficiency and specificity at the aphid salivary gland. J Gen Virol 78 (Pt 3):495–503.

9. Rochow WF. 1969. Biological properties of four isolates of barley yellow dwarf virus. Phytopathology 59:1580–1589.

10. Götz M, Popovski S, Kollenberg M, Gorovitz R, Brown JK, Cicero J, Czosnek H, Winter S, Ghanim M. 2012. Implication of *Bemisia tabaci* heat shock protein 70 in begomovirus - whitefly interactions. J Virol 86:13241–13252.

11. Kanakala S, Ghanim M. 2016. Implication of the whitefly *Bemisia tabaci* cyclophilin B protein in the transmission of *Tomato yellow leaf curl virus*. Front Plant Sci 7:1702.

12. Li C, Cox-Foster D, Gray SM, Gildow F. 2001. Vector specificity of *Barley yellow dwarf virus* (BYDV) transmission: identification of potential cellular receptors binding BYDV-MAV in the aphid, *Sitobion avenae*. Virology 286:125–133.

13. Liang Y, Gao X. 2017. The cuticle protein gene MPCP4 of *Myzus persicae* (Homoptera: Aphididae) plays a critical role in *Cucumber mosaic virus* acquisition. J Econ Entomol 110:848–853.

14. Liu W, Gray S, Huo Y, Li L, Wei T, Wang X. 2015. Proteomic analysis of interaction between a plant virus and its vector insect reveals new functions of hemipteran cuticular protein. Mol Cell Proteomics 14:2229–2242.

15. Webster CG, Pichon E, van Munster M, Monsion B, Deshoux M, Gargani D, Calvero F, Jimenez J, Moreno A, Krenz B, Thompson JR, Perry KL, Fereres A, Blanc S, Uzest M. 2018. Identification of plant virus receptor candidates in the stylets of their aphid vectors. J Virol 92:e00432–00418.

16. Xu Y, Wu J, Fu S, Li C, Zhu Z-R, Zhou X, Simon A. 2015. *Rice stripe tenuivirus* nonstructural protein 3 hijacks the 26S proteasome of the small brown planthopper via direct interaction with regulatory particle non-ATPase subunit 3. J Virol 89:4296–4310.

17. Wilson JR, DeBlasio SL, Alexander MM, Heck M. 2020. Looking through the lens of ‘omics technologies: Insights into the transmission of insect vector-borne plant viruses. Current Issues in Molecular Biology doi:10.21775/cimb.034.113:113-144.

18. Mulot M, Monsion B, Boissinot S, Rastegar M, Meyer S, Bochet N, Brault V. 2018. Transmission of *Turnip yellows virus* by *Myzus persicae* is reduced by feeding aphids on double-stranded RNA targeting the ephrin receptor protein. Front Microbiol 9:457.

19. Linz LB, Liu S, Chougule NP, Bonning BC. 2015. *In vitro* evidence supports membrane alanyl aminopeptidase N as a receptor for a plant virus in the pea aphid vector. J Virol 89:11203–11212.

20. Tamborindeguy C, Bereman MS, Igwe D, Deblasio S, Smith D, White F, MacCoss M, Gray S, Cilia M. 2013. Genomic and proteomic analysis of Schizaphis graminum reveals cyclophilin is involved in the transmission of Cereal yellow dwarf virus. PLoS One:e71620.

21. Woolston CJ, Covey SN, Penswick JR, Davies JW. 1983. Aphid transmission and a polypeptide are specified by a defined region of the *Cauliflower mosaic virus* genome. Gene 23:15–23.

22. Pirone TP, Perry KL. 2002. Aphids: non-persistent transmission. Adv Bot Res 36:1–19.

23. Badillo-Vargas IE, Chen Y, Martin KM, Rotenberg D, Whitfield AE. 2019. Discovery of novel thrips vector proteins that bind to the viral attachment protein of the plant bunyavirus tomato spotted wilt virus. J Virol 93:e00699–00619.

24. Dombrovsky A, Gollop N, Chen S, Chejanovsky N, Raccah B. 2007. *In vitro* association between the helper component-proteinase of *Zucchini yellow mosaic virus* and cuticle proteins of *Myzus persicae*. J Gen Virol 88:1602–1610.

25. Li S, Xiong R, Wang X, Zhou Y. 2011. Five proteins of *Laodelphax striatellus* are potentially involved in the interactions between rice stripe virus and vector. PLoS One 6:e26585.

26. Seddas P, Boissinot S, Strub JM, Van Dorsselaer A, Van Regenmortel MH, Pattus F. 2004. Rack-1, GAPDH3, and actin: proteins of *Myzus persicae* potentially involved in the transcytosis of beet western yellows virus particles in the aphid. Virology 325:399-412.

27. van den Heuvel JF, Verbeek M, van der Wilk F. 1994. Endosymbiotic bacteria associated with circulative transmission of potato leafroll virus by *Myzus persicae*. J Gen Virol 75 (Pt 10):2559–2565.

28. Li S, Li X, Zhou Y. 2018. Ribosomal protein L18 is an essential factor that promote *Rice stripe virus* accumulation in small brown planthopper. Virus Res 247:15–20.

29. Mar T, Liu W, Wang X. 2014. Proteomic analysis of interaction between P7-1 of *Southern rice black-streaked dwarf virus* and the insect vector reveals diverse insect proteins involved in successful transmission. J Proteomics 102.

30. Badillo-Vargas IE, Rotenberg D, Schneweis DJ, Hiromasa Y, Tomich JM, Whitfield AE. 2012. Proteomic analysis of *Frankliniella occidentalis* and differentially-expressed proteins in response to Tomato spotted wilt virus infection. J Virol doi:JVI.00285-12 [pii]10.1128/JVI.00285-12.

31. Brault V, Tanguy S, Reinbold C, Le Trionnaire G, Arneodo J, Jaubert-Possamai S, Guernec G, Tagu D. 2010. Transcriptomic analysis of intestinal genes following acquisition of pea enation mosaic virus by the pea aphid *Acyrthosiphon pisum*. J Gen Virol 91:802–808.

32. Cassone BJ, Wijeratne S, Michel AP, Stewart LR, Chen Y, Yan P, Redinbaugh MG. 2014. Virus-independent and common transcriptome responses of leafhopper vectors feeding on maize infected with semi-persistently and persistent propagatively transmitted viruses. BMC Genomics 15:133.

33. Shrestha A, Champagne DE, Culbreath AK, Rotenberg D, Whitfield AE, Srinivasan R. 2017. Transcriptome changes associated with *Tomato spotted wilt virus* infection in various life stages of its thrips vector, *Frankliniella fusca* (Hinds). J Gen Virol 98:2156–2170.

34. Wang L, Tang N, Gao X, Guo D, Chang Z, Fu Y, Akinyemi IA, Wu Q. 2016. Understanding the immune system architecture and transcriptome responses to *Southern rice black-streaked dwarf virus* in *Sogatella furcifera*. Sci Rep 6:36254.

35. Cilia M, Tamborindeguy C, Fish T, Howe K, Thannhauser TW, Gray S. 2011. Genetics coupled to quantitative intact proteomics links heritable aphid and endosymbiont protein expression to circulative polerovirus transmission. J Virol 85:2148–2166.

36. Papura D, Jacquot E, Dedryver CA, Luche S, Riault G, Bossis M, Rabilloud T. 2002. Two-dimensional electrophoresis of proteins discriminates aphid clones of Sitobion avenae differing in BYDV-PAV transmission. Arch Virol 147:1881–1898.

37. Yang X, Thannhauser TW, Burrows M, Cox-Foster D, Gildow FE, Gray SM. 2008. Coupling genetics and proteomics to identify aphid proteins associated with vector-specific transmission of polerovirus (luteoviridae). J Virol 82:291–299.

38. Mauck K, Bosque-Perez NA, Eigenbrode SD, De Moraes CM, Mescher MC. 2012. Transmission mechanisms shape pathogen effects on host-vector interactions: evidence from plant viruses. Functional Ecology 26:1162–1175.

39. Heck M, Brault V. 2018. Targeted disruption of aphid transmission: a vision for the management of crop diseases caused by *Luteoviridae* members. Curr Opin Virol 33:24–32.

40. DeBlasio SL, Johnson R, Mahoney J, Karasev A, Gray SM, MacCoss MJ, Cilia M. 2015. Insights into the polerovirus-plant interactome revealed by coimmunoprecipitation and mass spectrometry. Mol Plant Microbe Interact 28:467–481.

41. DeBlasio SL, Johnson R, Sweeney MM, Karasev A, Gray SM, MacCoss MJ, Cilia M. 2015. Potato leafroll virus structural proteins manipulate overlapping, yet distinct protein interaction networks during infection. Proteomics 15:2098–2112.

42. DeBlasio SL, Johnson RS, MacCoss MJ, Gray SM, Cilia M. 2016. Model system-guided protein interaction mapping for virus isolated from phloem tissue. J Proteome Res 15:4601–4611.

43. Conlon FL, Miteva Y, Kaltenbrun E, Waldron L, Greco TM, Cristea IM. 2012. Immunoisolation of protein complexes from Xenopus. Methods Mol Biol 917:369–390.

44. Rogstad SM, Sorkina T, Sorkin A, Wu CC. 2013. Improved precision of proteomic measurements in immunoprecipitation based purifications using relative quantitation. Anal Chem 85:4301–4306.

45. Choi H, Larsen B, Lin ZY, Breitkreutz A, Mellacheruvu D, Fermin D, Qin ZS, Tyers M, Gingras AC, Nesvizhskii AI. 2011. SAINT: probabilistic scoring of affinity purification-mass spectrometry data. Nat Methods 8:70–73.

46. Skarra DV, Goudreault M, Choi H, Mullin M, Nesvizhskii AI, Gingras AC, Honkanen RE. 2011. Label-free quantitative proteomics and SAINT analysis enable interactome mapping for the human Ser/Thr protein phosphatase 5. Proteomics 11:1508–1516.

47. Teo G, Liu G, Zhang J, Nesvizhskii AI, Gingras A-C, Choi H. 2014. SAINTexpress: Improvements and additional features in Significance Analysis of INTeractome software. Journal of Proteomics 100:37–43.

48. Old WM, Meyer-Arendt K, Aveline-Wolfe L, Pierce KG, Mendoza A, Sevinsky JR, Resing KA, Ahn NG. 2005. Comparison of label-free methods for quantifying human proteins by shotgun proteomics. Mol Cell Proteomics 4:1487–1502.

49. Budayeva HG, Cristea IM. 2014. A mass spectrometry view of stable and transient protein interactions. 806:263–282.

50. Ding T-B, Li J, Chen E-H, Niu J-Z, Chu D. 2019. Transcriptome profiling of the whitefly *Bemisia tabaci* MED in response to single infection of *Tomato yellow leaf curl virus*, *Tomato chlorosis virus*, and their co-infection. Frontiers in Physiology 10.

51. Pinheiro PV, Ghanim M, Alexander M, Rebelo A, Santos RS, Orsburn BC, Gray S, Cilia M. 2017. Host plants indirectly influence plant virus transmission by altering gut cysteine protease activity of aphid vectors. Mol Cell Proteomics 16:S230–S243.

52. Hu M, Crawford Simon A, Henstridge Darren C, Ng Ivan HW, Boey Esther JH, Xu Y, Febbraio Mark A, Jans David A, Bogoyevitch Marie A. 2013. p32 protein levels are integral to mitochondrial and endoplasmic reticulum morphology, cell metabolism and survival. Biochemical Journal 453:381–391.

53. McGee AM, Douglas DL, Liang Y, Hyder SM, Baines CP. 2014. The mitochondrial protein C1qbp promotes cell proliferation, migration and resistance to cell death. Cell Cycle 10:4119–4127.

54. Scully OJ, Yu Y, Salim A, Thike AA, Yip GW, Baeg GH, Tan PH, Matsumoto K, Bay BH. 2015. Complement component 1, q subcomponent binding protein is a marker for proliferation in breast cancer. Exp Biol Med (Maywood) 240:846–853.

55. Zhang X, Zhang F, Guo L, Wang Y, Zhang P, Wang R, Zhang N, Chen R. 2013. Interactome analysis reveals that C1QBP (complement component 1, q subcomponent binding protein) is associated with cancer cell chemotaxis and metastasis. Mol Cell Proteomics 12:3199–3209.

56. Hillman GA, Henry MF. 2019. The yeast protein Mam33 functions in the assembly of the mitochondrial ribosome. Journal of Biological Chemistry 294:9813–9829.

57. Muta J, Kang D, Kitajima S, Fugiwara T, Hamasaki N. p32 protein, a splicing factor 2-associated protein, is localized in mitochondrial matrix and is functionally important in maintaining oxidative phosphorylation. J Biol Chem 272:24363–24370.

58. Barna J, Dimén D, Puska G, Kovács D, Csikós V, Oláh S, Udvari EB, Pál G, Dobolyi Á. 2019. Complement component 1q subcomponent binding protein in the brain of the rat. Scientific Reports 9.

59. Soltys BJ, Kang D, Gupta RS. 2000. Localization of P32 protein (gC1q-R) in mitochondria and at specific extramitochondrial locations in normal tissues. Histochem Cell Biol 114:245–255.

60. Matthews DA, Russell WC. 1998. Adenovirus core protein V interacts with p32-- a protein which is associated with both the mitochondria and the nucleus. J Gen Virol 79 1677–1685.

61. van Leeuwen HC, O’Hare P. 2001. Retargeting of the mitochondrial protein p32/gC1Qr to a cytoplasmic compartment and the cell surface. J Cell Sci 114:2115–2123.

62. Wang Y, Finan JE, Middeldorp JM, Hayward SD. 1997. P32/TAP, a cellular protein that interacts with EBNA-1 of Epstein-Barr virus. Virology 236:18–29.

63. Ivanov S, Harrison MJ. 2014. A set of fluorescent protein-based markers expressed from constitutive and arbuscular mycorrhiza-inducible promoters to label organelles, membranes and cytoskeletal elements in *Medicago truncatula*. Plant J 80:1151–1163.

64. Negi S, Pandey S, Srinivasan SM, Mohammed A, Guda C. 2015. LocSigDB: a database of protein localization signals. Database 2015:bav003-bav003.

65. Yenugonda V, Nomura N, Kouznetsova V, Tsigelny I, Fogal V, Nurmemmedov E, Kesari S, Babic I. 2017. A novel small molecule inhibitor of p32 mitochondrial protein overexpressed in glioma. Journal of Translational Medicine 15.

66. Mellacheruvu D, Wright Z, Couzens AL, Lambert JP, St-Denis NA, Li T, Miteva YV, Hauri S, Sardiu ME, Low TY, Halim VA, Bagshaw RD, Hubner NC, Al-Hakim A, Bouchard A, Faubert D, Fermin D, Dunham WH, Goudreault M, Lin ZY, Badillo BG, Pawson T, Durocher D, Coulombe B, Aebersold R, Superti-Furga G, Colinge J, Heck AJ, Choi H, Gstaiger M, Mohammed S, Cristea IM, Bennett KL, Washburn MP, Raught B, Ewing RM, Gingras AC, Nesvizhskii AI. 2013. The CRAPome: a contaminant repository for affinity purification-mass spectrometry data. Nat Methods 10:730–736.

67. Cilia M, Johnson R, Sweeney M, DeBlasio SL, Bruce JE, MacCoss MJ, Gray SM. 2014. Evidence for lysine acetylation in the coat protein of a polerovirus. J Gen Virol 95:2321–2327.

68. Alexander MM, Mohr JP, DeBlasio SL, Chavez JD, Ziegler-Graff V, Brault V, Bruce JE, Heck MC. 2017. Insights in luteovirid structural biology guided by chemical cross-linking and high resolution mass spectrometry. Virus Res 241:42–52.

69. Patton MF, Bak A, Sayre JM, Heck ML, Casteel CL. 2019. A polerovirus, Potato leafroll virus, alters plant–vector interactions using three viral proteins. Plant, Cell & Environment 43:387–399.

70. Tahtouh M, Garcon-Bocquet A, Croq F, Vizioli J, Sautiere P, Van Camp C, Salzet M, Meillour PN, Pestel J, Lefebvre C. 2012. Interaction of *Hm*C1q with leech microglial cells: involvement of C1qBP-related molecule in the induction of cell chemotaxis. J Neuroinflammation 9:37.

71. Feichtinger RG, Oláhová M, Kishita Y, Garone C, Kremer LS, Yagi M, Uchiumi T, Jourdain AA, Thompson K, D’Souza AR, Kopajtich R, Alston CL, Koch J, Sperl W, Mastantuono E, Strom TM, Wortmann SB, Meitinger T, Pierre G, Chinnery PF, Chrzanowska-Lightowlers ZM, Lightowlers RN, DiMauro S, Calvo SE, Mootha VK, Moggio M, Sciacco M, Comi GP, Ronchi D, Murayama K, Ohtake A, Rebelo-Guiomar P, Kohda M, Kang D, Mayr JA, Taylor RW, Okazaki Y, Minczuk M, Prokisch H. 2017. Biallelic *C1QBP* mutations cause severe neonatal-, childhood-, or later-onset cardiomyopathy associated with combined respiratory-chain deficiencies. The American Journal of Human Genetics 101:525–538.

72. Jiang J, Zhang Y, Krainer AR, Xu R. 1999. Crystal structure of human p32, a doughnut-shaped acidic mitochondrial matrix protein. Proc Natl Acad Sci U S A 96:3572–3577.

73. Gotoh K, Morisaki T, Setoyama D, Sasaki K, Yagi M, Igami K, Mizuguchi S, Uchiumi T, Fukui Y, Kang D. 2018. Mitochondrial p32/C1qbp Is a Critical Regulator of Dendritic Cell Metabolism and Maturation. Cell Reports 25:1800–1815.e1804.

74. Krainer AR, Mayeda A, Kozak D, Binns G. 1991. Functional expression of cloned human splicing factor SF2: homology to RNA-binding proteins, Ul 70K, and Drosophila splicing regulators. Cell 66.

75. Son M, Diamond B, Santiago-Schwarz F. 2015. Fundamental role of C1q in autoimmunity and inflammation. Immunologic Research 63:101–106.

76. Xiao K, Wang Y, Chang Z, Lao Y, Chang DC. 2014. p32, a novel binding partner of Mcl-1, positively regulates mitochondrial Ca(2+) uptake and apoptosis. Biochem Biophys Res Commun 451:322–328.

77. Yu L, Zhang Z, Loewenstein PM, Desai K, Tang Q, Mao D, Symington JS, Green M. 1995. Molecular cloning and characterization of a cellular protein that interacts with the Human Immunodeficiency Virus Type 1 Tat transactivator and encodes a strong transcriptional activation domain. J Virol 69:3007–3016.

78. Zhang X, Zhang F, Guo L, Wang Y, Zhang P, Wang R, Zhang N, R. C. 2013. Interactome analysis reveals that C1QBP (complement component 1, q subcomponent binding protein) Is associated with cancer cell chemotaxis and metastasis. Molecular and Cellular Proteomics 12:3199–3209.

79. Ghebrehiwet B, Lim BL, Peerschke EI, Willis AC, KB R. 1994. Isolation, cDNA cloning, and overexpression of a 33-kD cell surface glycoprotein that binds to the globular “heads” of C1q. Journal of Experimental Medicine 179:1809-1821.

80. Dunkelberger JR, Song WC. 2010. Complement and its role in innate and adaptive immune responses. Cell Res 20:34–50.

81. Beatch M, Hobman TC. 2000. Rubella virus capsid associates with host cell protein p32 and localizes to mitochondria. J Virol 74:5569–5576.

82. Le Sage V, Cinti A, Valiente-Echeverría F, Mouland AJ. 2015. Proteomic analysis of HIV-1 Gag interacting partners using proximity-dependent biotinylation. Virology Journal 12.

83. Liu Z, Kato A, Oyama M, Kozuka-Hata H, Arii J, Kawaguchi Y, Sandri-Goldin RM. 2015. Role of host cell p32 in Herpes Simplex Virus 1 de-envelopment during viral nuclear egress. Journal of Virology 89:8982–8998.

84. Cohen S, Valm AM, Lippincott-Schwartz J. 2018. Interacting organelles. Current Opinion in Cell Biology 53:84–91.

85. Choi Y, Kwon Y-C, Kim S-I, Park J-M, Lee K-H, Ahn B-Y. 2008. A hantavirus causing hemorrhagic fever with renal syndrome requires gC1qR/p32 for efficient cell binding and infection. Virology 381:178–183.

86. Pednekar L, Valentino A, Ji Y, Tumma N, Valentino C, Kadoor A, Hosszu KK, Ramadass M, Kew RR, Kishore U, Peerschke EIB, Ghebrehiwet B. 2016. Identification of the gC1qR sites for the HIV-1 viral envelope protein gp41 and the HCV core protein: Implications in viral-specific pathogenesis and therapy. Molecular Immunology 74:18–26.

87. Song X, Yao Z, Yang J, Zhang Z, Deng Y, Li M, Ma C, Yang L, Gao X, Li W, Liu J, Wei L. 2016. HCV core protein binds to gC1qR to induce A20 expression and inhibit cytokine production through MAPKs and NF-κB signaling pathways. Oncotarget 7:33796–33808.

88. Xu L, Xiao N, Liu F, Ren H, Gu J. 2009. Inhibition of RIG-I and MDA5-dependent antiviral response by gC1qR at mitochondria. Proc Natl Acad Sci U S A 106:1530–1535.

89. Claus C, Chey S, Heinrich S, Reins M, Richardt B, Pinkert S, Fechner H, Gaunitz F, Schafer I, Seibel P, Liebert UG. 2011. Involvement of p32 and Microtubules in Alteration of Mitochondrial Functions by Rubella Virus. Journal of Virology 85:3881–3892.

90. de Graaf DC, Brunain M, Scharlaken B, Peiren N, Devreese B, Ebo DG, Stevens WJ, Desjardins CA, Werren JH, Jacobs FJ. 2010. Two novel proteins expressed by the venom glands ofApis melliferaandNasonia vitripennisshare an ancient C1q-like domain. Insect Molecular Biology 19:1–10.

91. Consortium TIAG. 2010. Genome sequence of the pea aphid Acyrthosiphon pisum. PLoS Biol 8:e1000313.

92. Pinheiro PV, Wilson JR, Xu Y, Zheng Y, Rebelo AR, Fattah-Hosseini S, Kruse A, Dos Silva RS, Xu Y, Kramer M, Giovannoni J, Fei Z, Gray S, Heck M. 2019. Plant viruses transmitted in two different modes produce differing effects on small RNA-mediated processes in their aphid vector. Phytobiomes Journal 3:71–81.

93. Burrows ME, Caillaud MC, Smith DM, Benson EC, Gildow FE, Gray SM. 2006. Genetic regulation of polerovirus and luteovirus transmission in the aphid *Schizaphis graminum*. Phytopathology 96:828–837.

94. Franco-Lara LF, McGeachy KD, Commandeur U, Martin RR, Mayo MA, Barker H. 1999. Transformation of tobacco and potato with cDNA encoding the full-length genome of potato leafroll virus: evidence for a novel virus distribution and host effects on virus multiplication. J Gen Virol 80 (Pt 11):2813–2822.

95. DeBlasio SL, Rebelo AR, Parks K, Gray S, Cilia M. 2018. Disruption of chloroplast function through downregulation of phytoene desaturase enhances the systemic accumulation of an aphid-borne, phloem-restricted virus. Mol Plant Microbe Interact doi:10.1094/MPMI-03-18-0057-R:doi: 10.1094/MPMI-1003-1018-0057-R.

96. DeBlasio S, Xu Y, Johnson R, Rebelo A, MacCoss M, Gray S, Heck M. 2018. The interaction dynamics of two Potato Leafroll Virus movement proteins affects their localization to the outer membranes of mitochondria and plastids. Viruses 10:585.

97. Peter KA, Liang D, Palukaitis P, Gray SM. 2008. Small deletions in the potato leafroll virus readthrough protein affect particle morphology, aphid transmission, virus movement and accumulation. J Gen Virol 89:2037–2045.

98. MacLean B, Tomazela DM, Shulman N, Chambers M, Finney GL, Frewen B, Kern R, Tabb DL, Liebler DC, MacCoss MJ. 2010. Skyline: an open source document editor for creating and analyzing targeted proteomics experiments. Bioinformatics 26:966–968.

99. Earley KW, Haag JR, Pontes O, Opper K, Juehne T, Song K, Pikaard CS. 2006. Gateway-compatible vectors for plant functional genomics and proteomics. Plant J 45:616–629.

